# Acquired Cross-resistance in Small Cell Lung Cancer due to Extrachromosomal DNA Amplification of *MYC* paralogs

**DOI:** 10.1101/2023.06.23.546278

**Authors:** Shreoshi Pal Choudhuri, Luc Girard, Jun Yi Stanley Lim, Jillian F. Wise, Braeden Freitas, Di Yang, Edmond Wong, Seth Hamilton, Victor D. Chien, Collin Gilbreath, Jun Zhong, Sarah Phat, David T. Myers, Camilla L. Christensen, Marcello Stanzione, Kwok-Kin Wong, Anna F. Farago, Catherine B. Meador, Nicholas J. Dyson, Michael S. Lawrence, Sihan Wu, Benjamin J. Drapkin

## Abstract

Small cell lung cancer (SCLC) presents as a highly chemosensitive malignancy but acquires cross-resistance after relapse. This transformation is nearly inevitable in patients but has been difficult to capture in laboratory models. Here we present a pre-clinical system that recapitulates acquired cross-resistance in SCLC, developed from 51 patient-derived xenografts (PDXs). Each model was tested for *in vivo* sensitivity to three clinical regimens: cisplatin plus etoposide, olaparib plus temozolomide, and topotecan. These functional profiles captured hallmark clinical features, such as the emergence of treatment-refractory disease after early relapse. Serially derived PDX models from the same patient revealed that cross-resistance was acquired through a *MYC* amplification on extrachromosomal DNA (ecDNA). Genomic and transcriptional profiles of the full PDX panel revealed that this was not unique to one patient, as *MYC* paralog amplifications on ecDNAs were recurrent among cross-resistant models derived from patients after relapse. We conclude that ecDNAs with *MYC* paralogs are recurrent drivers of cross-resistance in SCLC.

**SIGNIFICANCE:** SCLC is initially chemosensitive, but acquired cross-resistance renders this disease refractory to further treatment and ultimately fatal. The genomic drivers of this transformation are unknown. We use a population of PDX models to discover that amplifications of *MYC* paralogs on ecDNA are recurrent drivers of acquired cross-resistance in SCLC.

## INTRODUCTION

Small cell lung cancer (SCLC) remains one of the most common and deadly human cancers. Each year it afflicts 30,000-35,000 patients in the United States (200,000-250,000 patients worldwide) with a median survival of 7-11 months that has not improved significantly over the past 40 years (1-6). In 2012, the United States Congress designated SCLC as a “recalcitrant cancer” due to high incidence, poor prognosis, and stalled progress (7). More than 95% of SCLC present with disseminated disease (8,9), and even among the rare surgical cases only 40% are cured with resection alone (10). However, untreated SCLC is remarkably sensitive to DNA damage, and for these reasons, chemotherapy is recommended regardless of stage (11). First-line regimens combining etoposide with cisplatin (EP) or carboplatin (EC) yield overall response rates (ORRs) of 60-70%, rivaling the most effective oncogene-targeted therapies in non-small cell lung cancer (NSCLC) (12-15). A variety of agents can yield similar efficacy, including platinum-irinotecan combinations and cyclophosphamide-or methotrexate-containing regimens (16-19). This broad sensitivity to DNA damage distinguishes untreated SCLC from NSCLC, where responses are far less frequent (ORR 15-30%) (20,21).

For a small minority of patients, the initial response to chemotherapy can be extended for years with concurrent thoracic radiation or immune checkpoint blockade (22-26). However, for most patients, SCLC relapses within months as a far less manageable disease. Topotecan and lurbinectedin are approved second-line agents, but treatment responses are infrequent and brief (ORR 24% and 35%, and duration 3.3 and 5.3 months, respectively) (27,28). Other drugs have similar modest efficacy, including temozolomide (TMZ), amrubicin, gemcitabine, bendamustine, vinorelbine and taxanes (29,30). In addition to sharing similar efficacy, second-line agents share the same functional biomarker, the chemotherapy-free interval (CTFI) between conclusion of first-line chemotherapy and progression (30). “Platinum-resistant” SCLC (CTFI < 90 days) is unlikely to respond to any further chemotherapy (ORR 14% across regimens). In contrast, “platinum-sensitive” SCLC (CTFI > 90 days) may remain sensitive to first-line chemotherapy and can be re-challenged (ORR 49%) (31). However, this approach is only transiently effective, as patients with platinum-sensitive SCLC rarely survive more than 7-8 months regardless of regimen (31). This common trajectory suggests two categories of SCLC: cancers that remain sensitive to DNA damage and cancers that have acquired broad cross-resistance.

Acquired cross-resistance renders SCLC untreatable and rapidly fatal, as patients rarely receive more than 2-3 lines of therapy (32). Despite decades of research, the drivers of this transformation remain unknown. Scarcity of tumor samples has hindered research, as there is no clinical indication to re-biopsy SCLC after relapse. Furthermore, laboratory models that recapitulate clinical cross-resistance are not defined, as genetically engineered mouse models lack clinical histories, and cell lines derived before treatment or after relapse have similar chemosensitivity *in vitro* (33). Patient-derived xenograft (PDX) models may overcome these barriers. PDXs can be established after relapse from circulating tumor cells (CTCs), bypassing the need for re-biopsy (34), and maintaining genomic fidelity to their donor tumors (35). More importantly, PDX models recapitulate the responses of their corresponding patients to DNA-damaging regimens such as first-line EP or EC, and olaparib plus TMZ (OT), a promising experimental regimen for relapsed SCLC (35,36). This functional fidelity is necessary to capture acquired cross-resistance.

To investigate cross-resistance, we generated comprehensive clinical, molecular, and functional profiles of 51 SCLC PDX models. Each model was treated with three clinical regimens *in vivo* to quantify cross-resistance, and these values were compared with patient treatment histories to confirm clinical fidelity. A pair of serially derived PDXs from the same patient revealed that a focal amplification of *MYC* on extrachromosomal DNA (ecDNA) was acquired on treatment to confer cross-resistance. The full PDX panel demonstrated that this was not isolated to one patient, as ecDNA amplifications of *MYC, MYCN*, and *MYCL* were recurrent among cross-resistant models derived after relapse. We conclude that ecDNAs harboring *MYC* paralogs may drive acquired cross-resistance in SCLC.

## RESULTS

### Serial PDX models demonstrate acquired cross-resistance and high-level MYC expression

The clinical trajectory of SCLC is biphasic and defined by acquired cross-resistance (**Fig. 1A**). To discover genetic alterations that underlie this transition, we compared serial PDX models derived from the same patient, MGH1518, before first-line therapy with EC (MGH1518-1BX) and again after a durable response to second-line OT (MGH1518-3A) (**Fig. 1B**). These models recapitulated the patient’s clinical history, with MGH1518-3A acquiring resistance to both EP and OT (**Fig. 1C** and **Supplementary Figure S1A-C**; clinical efficacy of EC and EP are equivalent (12)). Resistance in this model extended beyond the regimens received by the patient to include topotecan, demonstrating acquired cross-resistance (**Fig. 1C**).

**Figure 1.**
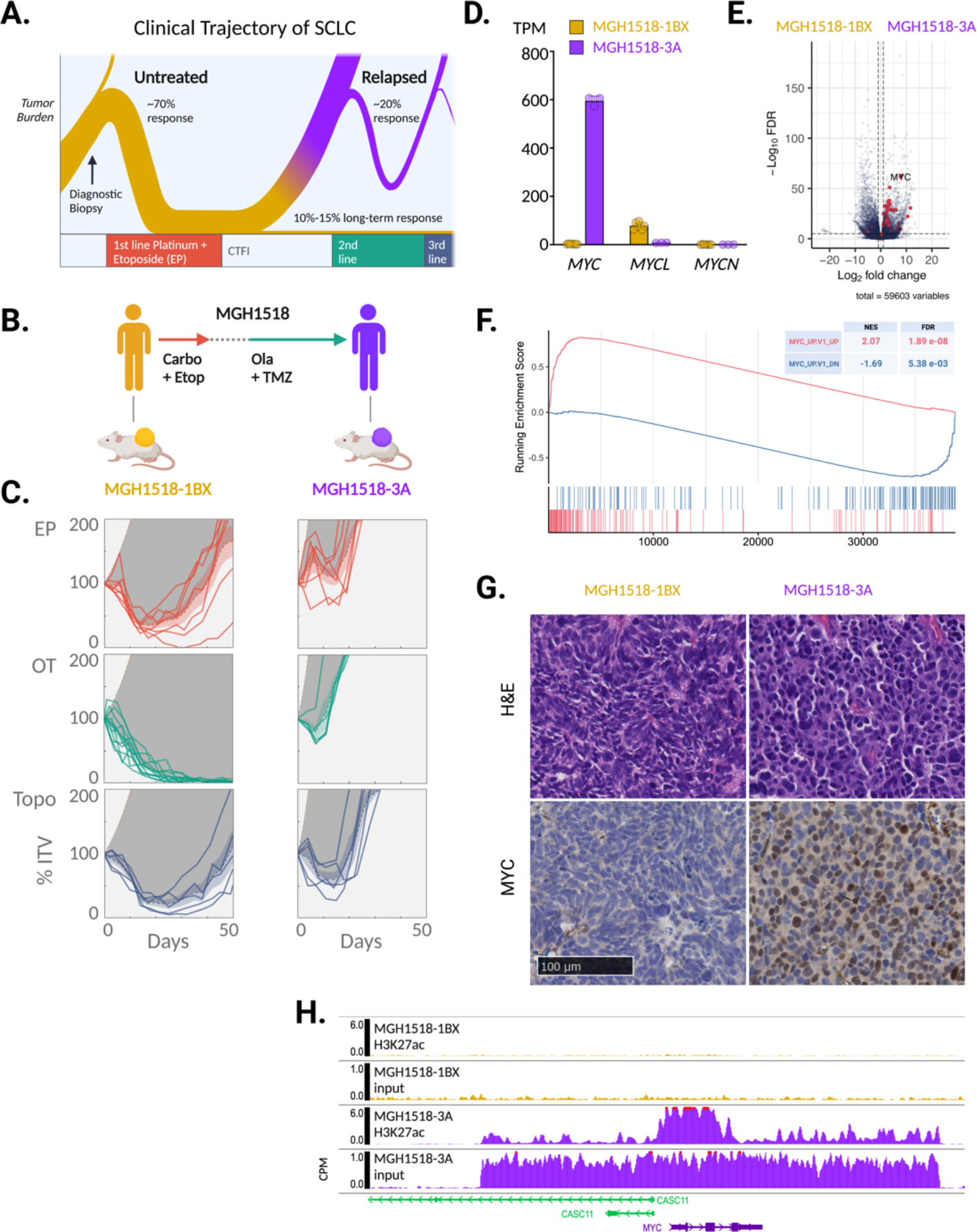
Serial PDX models of SCLC demonstrate cross-resistance and high-level MYC expression acquired after two lines of therapy. (**A**) The biphasic clinical trajectory of SCLC, from broad chemosensitivity to acquired cross-resistance. Ribbon thickness depicts proportion of patients. Purple = relapse. (**B**) Clinical treatment history of patient MGH1518. Prior to therapy, PDX model MGH1518-1BX was derived from core biopsy. Patient then received 5 cycles of EC, followed by 65 days off therapy and progression at first restaging. Second-line OT resulted in a durable partial response of 6.8 months. PDX model MGH1518-3A was derived after progression on OT. (**C**) PDX responses to EP, OT and topotecan (Topo), as described in detail in **Supplementary Figure S1A-C** for MGH1518-1BX treated with EP (top left). Solid color lines = tumor-volume (TV) curves for treated xenografts starting from initial tumor volume (ITV) of 300-600 mm^3^. Dashed color lines + color shading = average TV-curves ± 95% confidence interval (CI). Tan dashed lines + shading = untreated TV curves ± 95% CI from model growth coefficients calculated from 94 MGH1518-1BX xenografts and 60 MGH1518-3A xenografts. Dark gray shading = difference in area under treated and untreated average TV curves (DAUC). (**D**) PDX *MYC* paralog transcripts per million (TPM). Circles = replicate xenografts. Error bars = mean and SEM. (**E**) PDX differential gene expression. Red = genes upregulated upon overexpression of *MYC* in primary breast epithelial cells (MYC_UP.V1_UP). (**F**) Gene set enrichment plot in MGH1518-3A vs. MGH1518-1BX for genes upregulated and downregulated with MYC overexpression. (**G**) PDX morphology by hematoxylin and eosin stain, and MYC expression by immunohistochemistry. (**H**) PDX Histone H3K27ac ChIP-seq on chromosome 8 within 25 kb of *MYC* locus. Normalized input and H3K27ac peaks between models. Peak heights = counts per million mapped reads (CPM).

The underlying gene expression changes that might account for resistance were investigated through transcriptome sequencing. This revealed a striking upregulation of the basic helix-loop-helix transcription factor *MYC* in MGH1518-3A, with nearly absent expression in MGH1518-1BX and only modest expression of its paralog, *MYCL* (**Fig. 1D**). Upregulation of *MYC* was highly impactful, driving a global shift in gene expression (**Table 1**) that was led by changes characteristic of *MYC* induction (**Fig. 1E-F** and **Supplementary Fig. S1D**) (37). This *MYC*-driven transcriptomic change was reflected in the divergent cellular morphologies of the tumors (**Fig. 1G**). MGH1518-1BX demonstrated features of classic SCLC, with sheets of fusiform cells with scant cytoplasm, nuclear molding, and inconspicuous nucleoli. In contrast, MGH1518-3A demonstrated the variant morphology originally associated with *MYC* amplification, with more abundant cytoplasm, round nuclei, and occasional nucleoli (38).

Increased *MYC* transcription resulted in elevated MYC protein levels and nuclear concentration, as measured by immunoblot and immunohistochemistry (**Fig. 1G** and **Supplementary Fig. S1E**). MYC staining was surprisingly heterogeneous, with nuclear signal nearly absent in some cells and intensely positive in others. To discover epigenetic changes that might account for variable high-level *MYC* induction, we profiled transcriptionally active chromatin through histone H3K27 acetylation chromatin immunoprecipitation and sequencing (H3K27ac ChIP-seq). This revealed a massively increased H3K27ac mark on the *MYC* locus in MGH1518-3A as compared to MGH1518-1BX, in agreement with their sharp difference in *MYC* transcription levels (**Fig. 1H**). More importantly, input chromatin from MGH1518-3A suggested a DNA amplification for *MYC* and the surrounding region (**Fig. 1H**). Together, these results suggested a focal amplification of *MYC* resulting in high-level but heterogeneous gene expression. This could be explained by either sub-clonal amplification or by amplification on circular extrachromosomal DNA (ecDNA), which is known to drive tumor heterogeneity (39).

### High-level MYC amplification on ecDNA was acquired after the start of second-line therapy

To investigate the focal *MYC* amplification further, MGH1518-1BX and MGH1518-3A were compared by whole genome sequencing (WGS). Most copy number variations were conserved between the models, but MGH1518-3A demonstrated a private 58-copy amplification of 2.09 Mb on chromosome 8 containing the *MYC* locus (**Fig. 2A** and **Supplementary Fig. S2A**). Reconstruction of the *MYC* amplicon with the AmpliconArchitect computational tool revealed an ecDNA composed of 22 fragments (**Fig. 2B** and **Supplementary Fig. S2B**) (40,41). The ecDNA was confirmed by *MYC* fluorescence in-situ hybridization (FISH) in metaphase cells, which showed multiple dispersed extrachromosomal foci (**Fig. 2C**). High-level focal amplifications of oncogenes and resistance factors on ecDNAs are common across cancer subtypes, including SCLC (42-44). Asymmetric segregation and positive selection of ecDNAs at mitosis not only allow them to reach extremely high copies per cell, but also generate intratumoral copy number heterogeneity (39), explaining the heterogeneity of MYC expression observed in MGH1518-3A cells (**Fig. 1G**).

**Figure 2.**
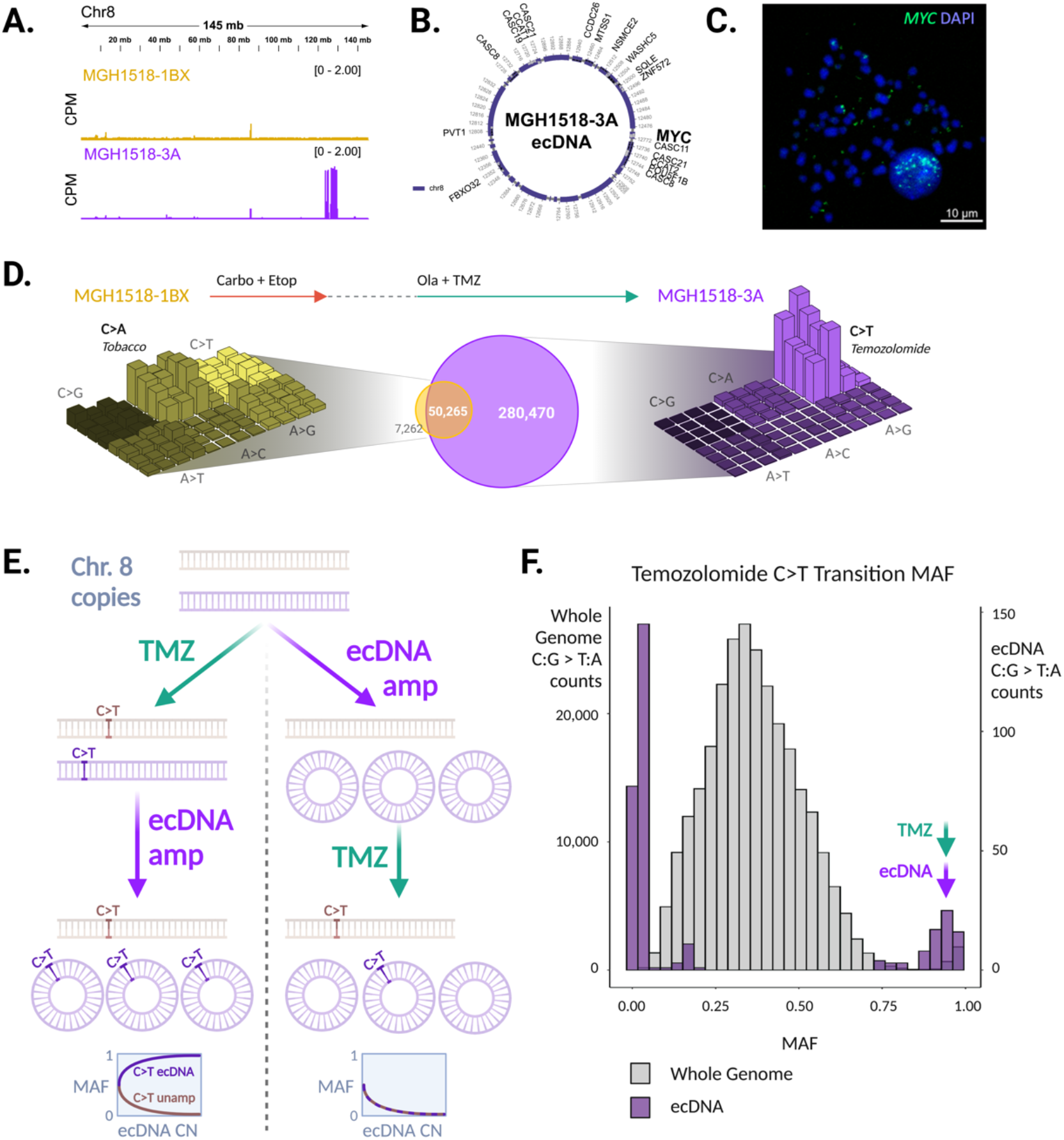
Serial PDX models demonstrate *MYC* amplification on ecDNA acquired after the start of second-line therapy. (**A**) Copy number variation on chromosome 8 demonstrated focal amplification of discontinuous segments including *MYC* in MGH1518-3A but not MGH1518-1BX. Peak heights = counts per million mapped reads (CPM). (**B**) AmpliconArchitect reconstruction of circular ecDNA containing *MYC*. (**C**) FISH using probes for *MYC* in metaphase cells from MGH1518-3A xenograft following dissociation. *MYC* foci apart from DAPI-stained chromosomes confirm presence of ecDNAs. (**D**) Center: Venn diagram compares the shared and unique mutations in serial models to reveal >280,000 new mutations in MGH1518-3A. Left and Right: three-dimensional bar plots (also called “Lego plots”) representing mutational signatures in a three-base context for each model (three-base key in **Figure S2C**). This reveals that following therapy with EP and OT, the shared G:C > T:A (C>A) transversions associated with tobacco smoking in MGH1518-1BX (left, yellow) have been overshadowed by unique C:G > T:A (C>T) transitions associated with TMZ in MGH1518-3A (right, purple). (**E**) Models depicting ecDNA formation either after (left) or before (right) massive influx of C>T mutations with OT treatment. If TMZ causes a C>T mutation before this mutation is amplified on the ecDNA (left), then the mutant allele frequency (MAF) of that mutation will increase with ecDNA copy number (CN), whereas conversely if TMZ causes a C>T mutation on the un-amplified allele then its MAF will decrease. If the ecDNA amplification occurs before TMZ (right), then C>T mutation MAFs will be inversely proportional to ecDNA CN regardless of location. (**F**) Distribution of C>T MAFs across the MGH1518-3A genome (grey) and on ec*MYC* (purple), in 30 bins ranging from 0 to 1. Bimodal distribution is consistent with ecDNA amplification after start of OT, as predicted by the left-side model in **Figure 2E**.

The ecDNA *MYC* amplification (ec*MYC*) was detected in MGH1518-3A but not in MGH1518-1BX (**Fig. 2A**). This may be due to emergence after MGH1518-1BX was derived, but alternative explanations are possible, such as the presence of ec*MYC* in a small subclone omitted from the cells that originated MGH1518-1BX. We reasoned that therapy-induced mutation signatures could be exploited to distinguish these possibilities. They consist of thousands of mutations across the genome, and comparison of the serial models revealed their emergence after therapy without ambiguity. MGH1518-3A was derived after the patient received EC and OT, and its tumor mutational burden increased by more than 6-fold (**Fig. 2D** and (45)). These new mutations were almost exclusively C>T transitions characteristic of TMZ (SBS11 in COSMIC), which became so abundant after relapse that they supplanted tobacco-induced C>A transversions (SBS4 in COSMIC) as the predominant mutation signature (**Fig. 2D** and **Supplementary Fig. S2C**) (46).

We analyzed the mutant allele frequencies (MAFs) of TMZ-induced C>T transitions to determine the timing of ec*MYC* amplification relative to the start of OT. Nearly every C>T mutation present in MGH1518-3A was absent from MGH1518-1BX, reflecting the TMZ-induced hypermutation described in glioblastoma (**Fig. 2D**) (47). If the ec*MYC* amplification were present before the start of OT, then the MAFs for C>T mutations within the ec*MYC* region would be diluted with increasing copy number (inversely proportional) (**Fig. 2E**). Even a low-level amplification would limit these MAFs to less than 0.5. In contrast, if ec*MYC* amplification occurred after starting OT, then the C>T MAFs within the ec*MYC* region would diverge with increasing copy number, depending on where they occurred (**Fig. 2E**). Mutations in the segments that later formed the ec*MYC* would be co-amplified, with their MAFs increasing with copy number, whereas mutations on the un-amplified allele would be diluted, generating a bimodal distribution. We compared the MAF distributions of C>T transitions on ec*MYC* with the whole-genome distribution. As expected, the distribution of C>T MAFs across the genome was unimodal, with a median of 0.37 reflecting overall near-triploidy (**Fig. 2F** and **Supplementary Figure S2A**). In contrast, the distribution of C>T MAFs within the ec*MYC* region was bimodal, as predicted if TMZ hypermutation occurred before ecDNA formation (**Fig. 2F** and **Table 2**). The difference between the peaks, with MAFs most frequent below 0.07 and above 0.90, reflects the high copy number of ec*MYC*.

We concluded that the ec*MYC* amplification was acquired in patient MGH1518’s SCLC after the start of second-line OT and drove the extreme changes in gene expression that accompanied acquired cross-resistance. This discovery raised two key questions. The first was whether ecDNA amplifications are frequently associated with cross-resistance in SCLC, with the caveat that this would require an experimental system to quantify cross-resistance. The second was whether ecDNA can drive cross-resistance. Both questions are addressed below.

### PDX models of SCLC recapitulate acquired cross-resistance to DNA-damaging agents

To understand whether ecDNA amplification is a generalized driver of cross-resistance, we developed a PDX-based experimental system to quantify therapy responses in SCLC and to investigate their association with molecular features. A panel of 51 PDX models were derived from 43 SCLC patients, 19 models before treatment and 32 after at least one line of therapy. For each model, the clinical history was annotated retrospectively to catalog all systemic therapies administered before and after model derivation, whether these resulted in clinical benefit, and the duration of therapies (**Fig. 3A**). For all but three patients (MGH1517, MGH1535, and MGH1586), a three-year window captured these clinical histories.

**Figure 3.**
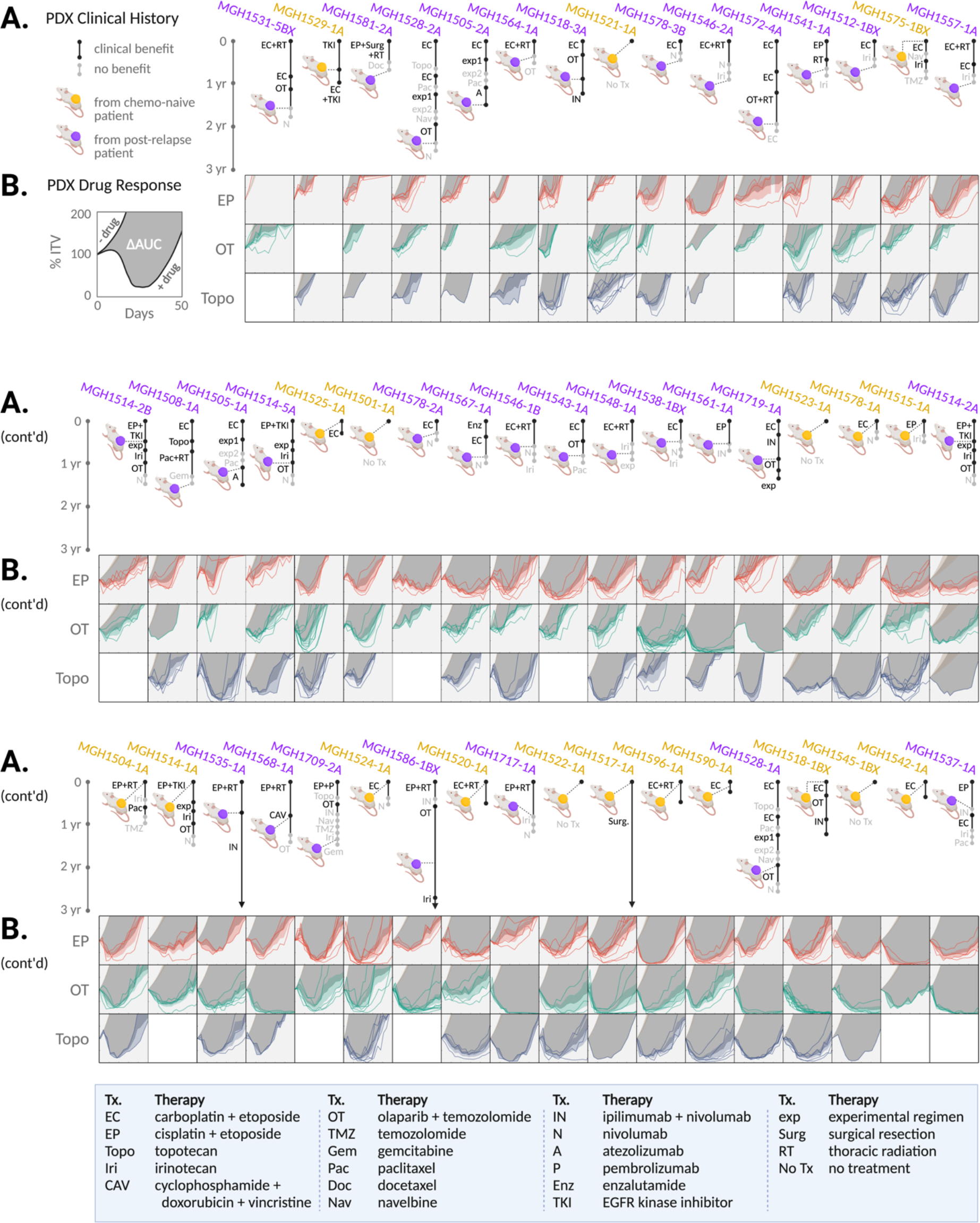
Clinical and functional annotation of PDX panel to measure acquired cross-resistance in SCLC. (**A**) Patient treatment histories before and after model derivation. Xenograft color denotes models derived before (yellow) or after (purple) first-line chemotherapy. Segment length = therapy duration. Segment shade = early progression (gray) vs. disease stabilization or regression (black). Therapy abbreviation key below. (**B**) PDX responses *in vivo* to EP, OT and topotecan. TV curves for individual treated xenografts, average TV curves for models with or without treatment ± 95% CI, and DAUC as depicted for MGH1518 models in **Figure 1C** and **Supplementary Figure S1A-C**. PDX models arranged by increasing chemosensitivity (ΔAUC^avg^, **Fig. 4D**) from top left to bottom right.

To measure cross-resistance directly, each model was treated *in vivo* with two to three clinical regimens that induce DNA damage through distinct mechanisms: EP, OT, and topotecan. Treatment was initiated at a large tumor volume (300-600 mm^3^) to measure regression precisely. Tumor-volume curves were compiled for each regimen, and an area-under-the-curve (AUC) bounded by 50 days and 200% initial tumor volume (ITV) was calculated for each xenograft (**Supplementary Fig. S1B**). The AUC metric was less subject to single-measurement variation than maximum regression or time-to-volume endpoints that are typically reported. Multiple xenografts were treated for each model and regimen, resulting in 678 tumor-volume curves (**Fig. 3B**). To control for intrinsic growth rate, the untreated AUC for each model was calculated from 1,184 control xenografts (**Supplementary Fig. S3A**). Model drug effect was calculated as the difference in mean AUC between drug-treated and control xenografts (ΔAUC) (**Supplementary Fig. S1A-C**). The *in vivo* regimens for EP, OT, and topotecan were calibrated such that the means and distributions of their ΔAUC values across the PDX panel were indistinguishable (**Supplementary Fig. S3B**). Furthermore, there was a strong correlation between ΔAUC values for each pair of regimens, with the correlation between EP and topotecan particularly strong (Pearson r = 0.66, p = 4.9e-6) (**Fig. 4A-C**). Therefore, we use a simple average of ΔAUC as a metric for modeling cross-resistance (ΔAUC^avg^) (**Fig. 4D** and **Table 3**).

**Figure 4.**
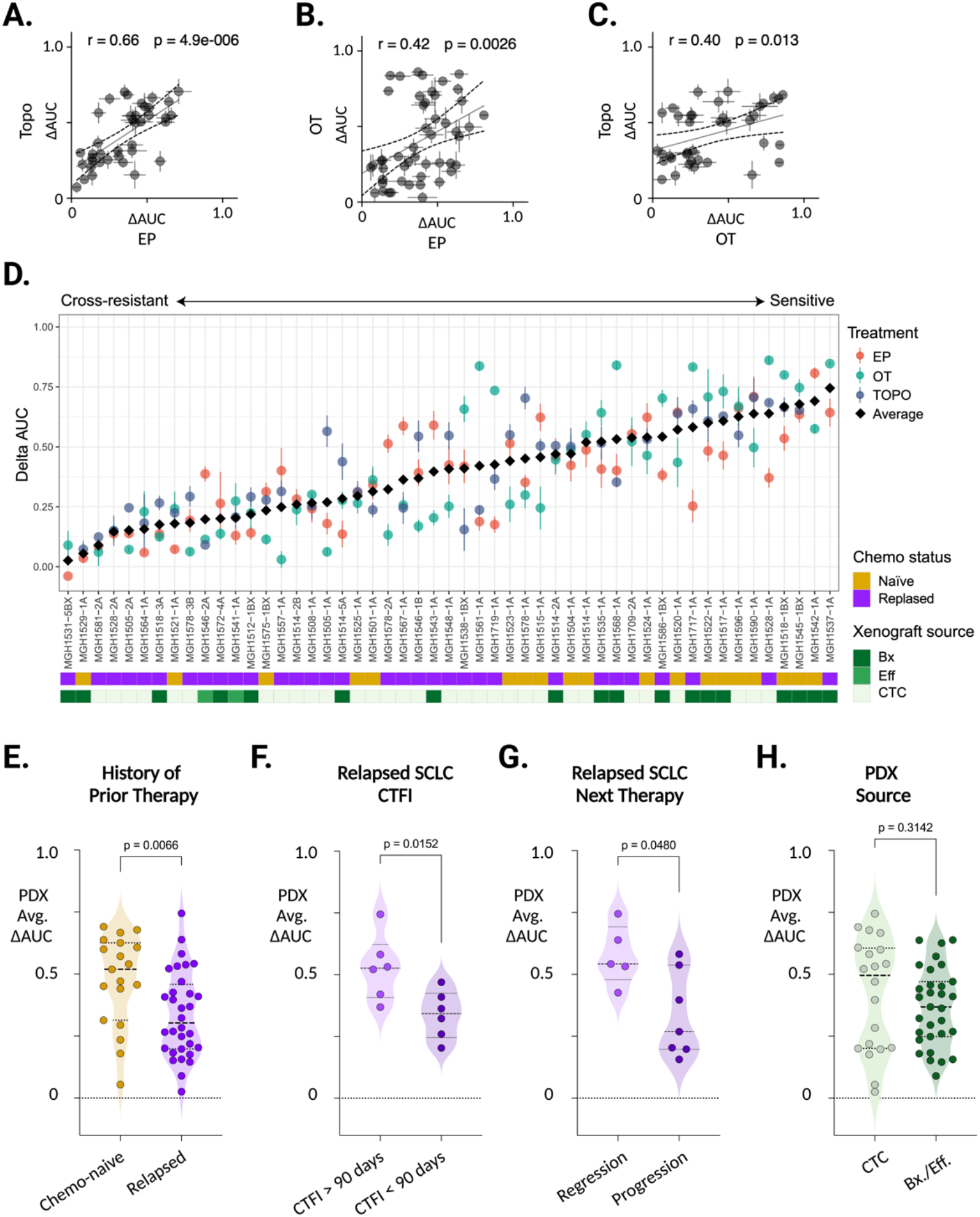
SCLC PDX models capture cross-resistance with high fidelity to the clinical treatment histories of their corresponding patients. (**A-C**) For each PDX model, pairwise comparison of ΔAUC for each regimen. Topotecan (Topo) vs. EP (**A**), OT vs. EP (**B**), and Topo vs. OT (**C**). Error bars = SEM of ΔAUC for each regimen. r = Pearson correlation, and p = probability that correlation is significant. Solid line = linear regression, dashed = 95% CI of regression. (**D**) Cross-resistance metric across the PDX panel. Models arranged left-to-right by average of ΔAUC for EP, OT, and Topo (ΔAUC^avg^), with annotation of clinical treatment history and PDX source below. (**E-G**) Performance of PDX ΔAUC^avg^ as a metric of clinical cross-resistance by comparison with features of patient treatment histories. Mann-Whitney test p-values for pairwise comparisons of ΔAUC^avg^: PDX models derived from (**E**) patients with untreated vs relapsed SCLC, (**F**) patients after first-line therapy with CTFI >90 days (platinum-sensitive) vs. <90 days (platinum-resistant), (**G**) patients with relapsed SCLC who responded vs. did not respond to next line of chemotherapy after model derivation. (**H**) ΔAUC^avg^ of PDX models derived from CTCs vs. biopsies or effusion specimens, compared as in **Figure 4E-G**.

When models were ordered by ΔAUC^avg^, cross-resistance between EP, OT and topotecan became apparent (**Fig. 3B**). To determine whether PDX ΔAUC^avg^ recapitulated clinical features of cross-resistance in SCLC, we compared this metric against known clinical determinants of treatment efficacy. Unlike established SCLC cell lines (33), PDX models derived from relapsed patients are significantly more cross-resistant than models from chemo-naïve patients (**Fig. 4E**). The overlap of ΔAUC^avg^ between these cohorts reasonably approximates the proportions of chemo-resistance among untreated SCLC and chemo-sensitivity in the relapsed population. To further test the fidelity of PDX model drug responses to clinical histories, we compared models by CTFI and by whether the patient’s next regimen was effective. Models derived after the first relapse from patients with a CTFI > 90 days (“platinum-sensitive”) had significantly higher ΔAUC^avg^ values that were comparable to models derived from untreated patients (**Fig. 4F**). Similarly, models derived from relapsed patients who responded to their next line of therapy were more chemosensitive than models derived just before an ineffective therapy (**Fig. 4G**). Clinical history distinctions were not confounded by PDX source, as there was no difference in ΔAUC^avg^ between CTC- and biopsy/effusion-derived models (**Fig. 4H**).

These results demonstrate a remarkable degree of fidelity for PDX model drug responses to the clinical histories of their patients. They also demonstrate the value of functional testing, as clinical history alone does not faithfully predict cross-resistance. Based on the history of relapse alone, highly sensitive models derived after a prolonged CTFI or from patients who responded to further chemotherapy would be grouped with cross-resistant SCLC, and would confound the search for candidate drivers of resistance (**Supplementary Fig. S3C**). We conclude that the PDX panel with both clinical and functional annotation is suitable for investigating the molecular features of cross-resistant SCLC.

### Cross-resistant PDX models of relapsed SCLC are enriched for *MYC*-family gene amplifications on ecDNAs (ec*MYC/N/L*)

To address whether ecDNA amplifications are recurrent with acquired cross-resistance in SCLC, we generated comprehensive genomic and transcriptomic profiles of each model by WGS and paired-end RNA sequencing. AmpliconArchitect reconstruction from WGS revealed ecDNAs in 23/51 PDX models (45%) (**Fig. 5A** and **Supplementary Fig. S4-5**), with *MYC* and its paralogs *MYCN* and *MYCL* as the most frequently amplified genes (ec*MYC/N/L*) (**Supplementary Fig. S6A**). All but one ec*MYC/N/L*+ model was derived after relapse (7/8), whereas ecDNAs lacking *MYC* paralogs were evenly distributed (5/19 chemo-naïve models vs. 10/32 post-relapse) (**Fig. 5A** and **Supplementary Fig. S4-5**). Models with ec*MYC/N/L* were significantly more cross-resistant than models that lacked ecDNAs, whereas there was no significant association between non-*MYC* ecDNAs and cross-resistance (**Fig. 5B**). The single chemo-naïve model harboring ec*MYCL*, MGH1501-1A, was also relatively cross-resistant (**Fig. 5A** and **Supplementary Fig. S4C-D**), suggesting that *de novo* ec*MYC/N/L* amplifications may underly poor responses to first-line therapy. The presence of ec*MYCL* in MGH1501-1A was confirmed by *MYCL* FISH in metaphase tumor cells (**Supplementary Fig. S4E**). It is unknown whether patient MGH1501 would have had primary refractory disease, as he passed away due to critical illness without receiving first-line therapy.

**Figure 5.**
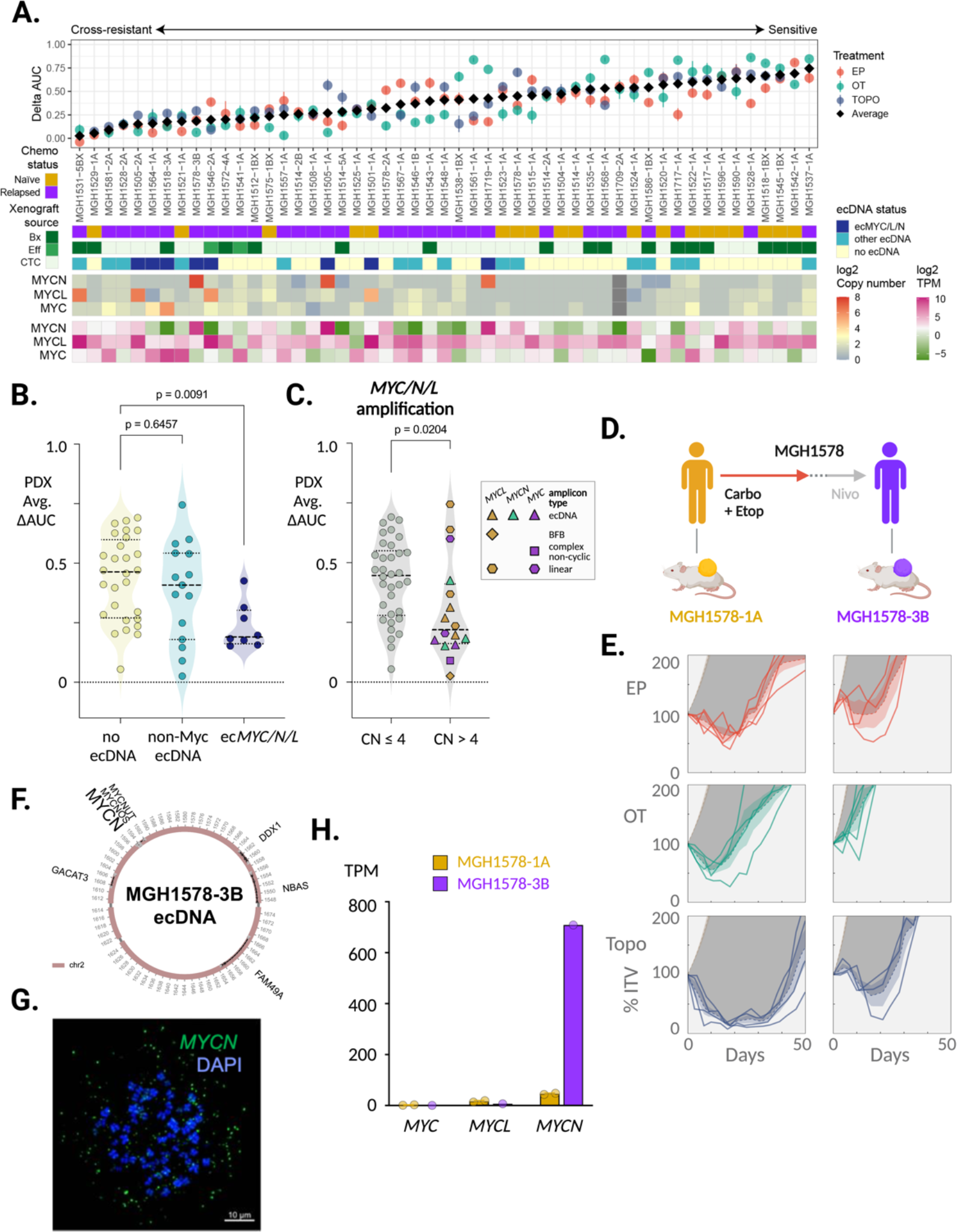
ecDNA amplifications of *MYC* paralogs are recurrent in cross-resistant PDX models derived from relapsed SCLC patients. (**A**) Integrated genomic landscape of *MYC* paralogs across SCLC PDX panel. Models arranged left-to-right by cross-resistance (ΔAUC^avg^) with annotated clinical history (chemo-naïve vs. post-relapse) and PDX source as in **Figure 4D**. Below: detection of ecDNA by AmpliconArchitect with or without *MYC* paralog (ec*MYC/N/L*) and copy numbers and transcript levels of *MYC* paralogs. (**B**) Comparison of ΔAUC^avg^ between models without ecDNAs and models with ecDNAs with or without *MYC* paralogs. Kruskal-Wallis test p-values for comparison with no-ecDNA control column. (**C**) Comparison of ΔAUC^avg^ between models with or without *MYC* paralog amplifications (CN > 4), with annotation of gene and amplification structure. Mann-Whitney test p-values. (**D**) Clinical treatment history of patient MGH1578. Prior to therapy, PDX model MGH1578-1A was derived from CTCs. Patient then received five cycles of EC reduced for cytopenias, followed by 14 days off therapy before progression. Second-line nivolumab was ineffective, then MGH1578-3B was derived from CTCs. (**E**) MGH1578 serial PDX responses to EP, OT and Topo as in **Figure 1C**, demonstrating acquired cross-resistance (**F-G**) AmpliconArchitect reconstruction of ec*MYCN* detected in MGH1578-3B but not MGH1578-1A, and confirmation with FISH of probes for *MYCN* in metaphase cells, as in **Figure 2B-C**. (**H**) MGH1578 PDX *MYC* paralog transcripts per million (TPM). Circles = replicate xenografts. Error bars = mean and SEM, as in **Figure 1D**.

The relationships between cross-resistance and *MYC* paralog copy number, expression level and amplicon type were analyzed. PDX models with a *MYC* paralog amplification (CN > 4) were more resistant than models with copy numbers within the range of genome duplication (CN ≤ 4), with ecDNA amplifications among the most resistant (**Fig. 5C**). ecDNAs accounted for all but one extreme amplification, a breakage-fusion-bridge (BFB) structure that resulted in 88 copies of *MYCL* in the highly resistant model MGH1531-5B (**Supplementary Fig. S6B**). Regression analysis of copy number, transcript level, and presence on an ecDNA revealed that all three features were significantly associated with cross-resistance (low ΔAUC^avg^), and ec*MYC/N/L* was associated with resistance to each regimen (EP, OT, topotecan) (**Supplementary Fig. S6C** and **Table 3**). We conclude that ecDNA formation is the most common mechanism for extreme amplification and expression of *MYC* paralogs, and that these structures are recurrent among cross-resistant SCLC tumors from relapsed patients.

The association of ec*MYC/N/L* with acquired cross-resistance was bolstered by a second set of serial models from patient MGH1578 (**Fig. 5D**). MGH1578-1A was derived from CTCs before first-line EC. After initial tumor regression, early relapse (CTFI < 90 days) and progression through second-line nivolumab, MGH1578-3B was derived from CTCs. MGH1578-3B demonstrated acquired resistance to EP as well as cross-resistance to OT and topotecan (**Fig. 5E**). WGS analysis revealed a massive 186-copy *MYCN* amplification in MGH1578-3B that was not present in the pre-treatment model (**Supplementary Fig. S7A**). AmpliconArchitect reconstruction revealed an ecDNA amplification confirmed by *MYCN* FISH in metaphase tumor cells (**Fig. 5F-G** and **Supplementary Fig. S7B**). Transcriptome comparison between MGH1578-1A and MGH1578-3B revealed a 15.5-fold increase in *MYCN* expression (**Fig. 5H**) that was reflected in differential expression of *MYC*-regulated gene sets (**Supplementary Fig. S7C-E**), as was observed in the MGH1518 serial models. In summary, ec*MYC/N/L* amplifications were enriched in cross-resistant PDX models of relapsed SCLC. For two sets of serial models, MGH1518 and MGH1578, the ecDNA was only detected after relapse and resistance. For MGH1518, mutation signature analysis demonstrated directly that the ec*MYC* amplification emerged after the start of second-line therapy (**Fig. 2D-F**). Together these results suggest that ec*MYC/N/L* amplifications are recurrent genomic alterations in relapsed cross-resistant SCLC.

### High-level ec*MYC* amplification confers resistance to DNA damage

Unlike chromosomes, ecDNAs lack centromeres to mediate spindle attachment during mitosis, instead segregating randomly to generate a binomial distribution in daughter cells (48). This natural heterogeneity provides an opportunity to investigate the effects of copy number variation in adjacent cells within the same tumor, without further genetic manipulation. We exploited this powerful tool to address whether ec*MYC* drives cross-resistance in SCLC. We asked whether chemotherapy induced less DNA damage in cells that inherit higher ec*MYC* copy numbers, and whether these cells are enriched in the tumor following treatment (**Fig. 6A**).

**Figure 6:**
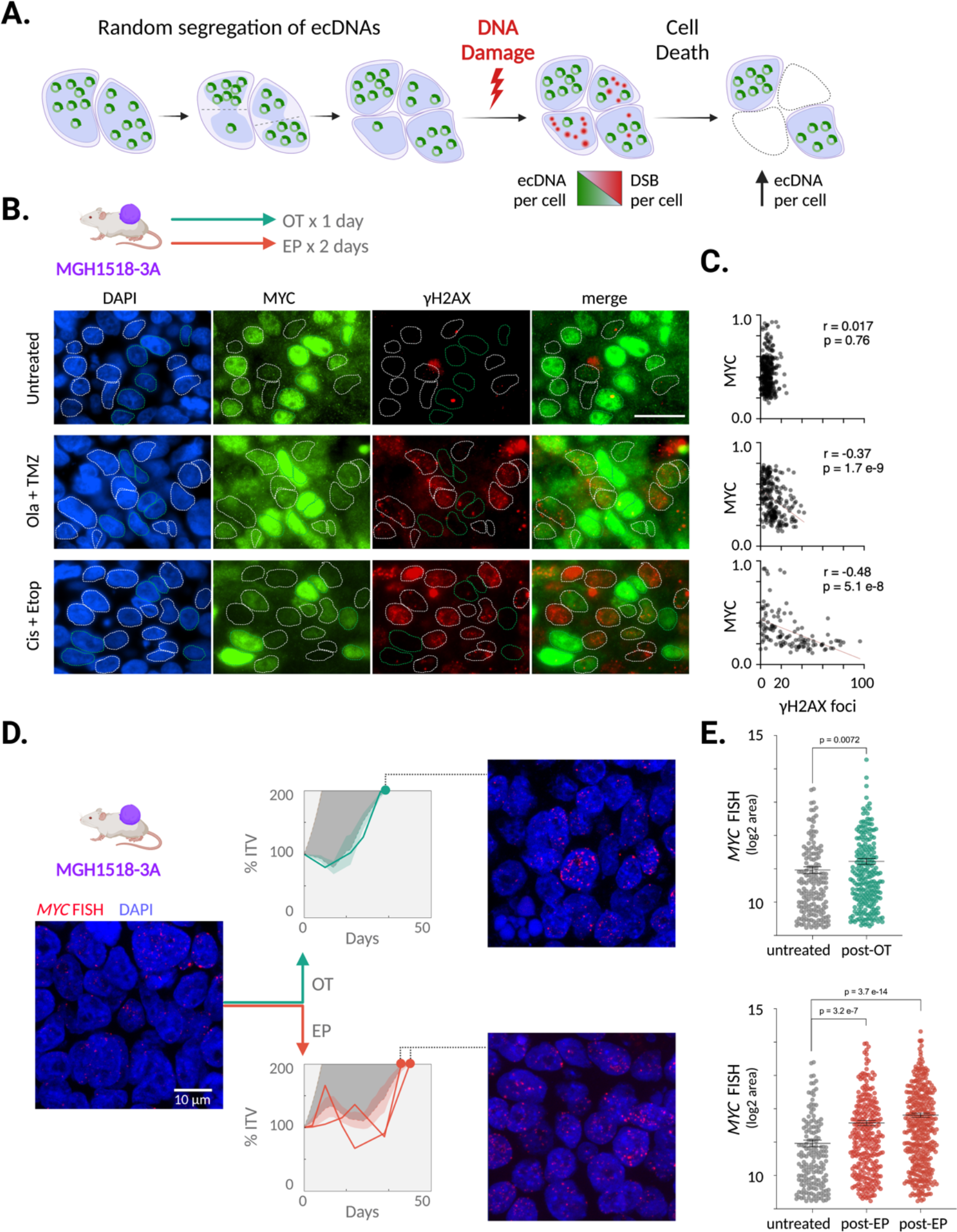
ec*MYC* confers dose-dependent resistance to DNA damaging agents. (**A**) Schema to determine whether ec*MYC* confers cross-resistance. Daughter cells inherit variable numbers of ecDNAs due to random segregation during mitosis. This generates natural copy-number heterogeneity that can be exploited experimentally to measure the effects of ecDNA dosage. Immediately after drug exposure, single-cell measurements of ec*MYC* and γH2AX foci are obtained to determine whether increasing ec*MYC* reduces DNA damage induction. Following xenograft progression, the distributions of ec*MYC* copy numbers in treated tumors are compared with untreated tumors to determine whether *ecMYC* promotes DNA damage survival. (**B-C**) MGH1518-3A xenografts (mean 58 copies ec*MYC* per cell) received either no treatment, a single day of OT or two days of EP and then were resected within 24 hours. (**B**) Dual immunofluorescence of MYC and γH2AX, with DAPI nuclear stain. (**C**) Normalized cellular MYC immunofluorescence vs. number of γH2AX. r = Pearson correlation coefficient, p = significance of correlation. (**D-E**) MGH1518-3A xenografts received full regimens of OT or EP and were resected after progression to 200% ITV. (**D**) FISH of probes for *MYC* in interphase cells with or without treatment. TV curves for treated xenografts superimposed on average TV curves for MGH1518-3A treated with OT or EP. (**E**) Measurement of *MYC* FISH area per cell in untreated and treated xenografts. Comparison of distributions, and non-parametric test p-values.

MGH1518-3A xenografts were treated with short courses of EP or OT and resected within 24 hours of the final dose to measure the effect of ec*MYC* copy number on DNA damage (**Fig. 6B**). Discrete nuclear foci of phosphorylated H2AX (γH2AX) form rapidly at double-stranded breaks, and their levels provide an indirect measure of the generation and repair of these lesions. Similarly, MYC expression levels are heterogeneous and may serve as a proxy for ec*MYC* copy number (**Fig. 1G** and **Fig. 6B**). In untreated tumors, γH2AX foci were rare and uncorrelated with MYC expression, but both EP and OT induced abundant γH2AX foci within 24 hours of the final dose. Strikingly, this damage was concentrated in the MYC^low^ cells, and almost all cells with the highest MYC expression were spared (**Fig. 6B**). This anticorrelation between γH2AX foci and MYC expression was both strong and highly significant, regardless of regimen (**Fig. 6C**), demonstrating that the cross-resistant cells inherited the highest ec*MYC* levels.

Although MGH1518-3A xenografts are highly cross-resistant compared with other PDX models (**Fig. 3** and **Fig. 4D**), their growth stalled transiently on treatment, with occasional regressions (**Fig. 1C**). Given the intratumoral heterogeneity of cancers with randomly segregating ecDNAs, we reasoned that this pause could represent a selection process for cells with an optimal ec*MYC* copy number (**Fig. 6B**). To test directly whether ec*MYC* confers a survival advantage, we performed *MYC* FISH on MGH1518-3A xenografts resected after regrowth following OT and EP (**Fig. 6D**). Following treatment, the ec*MYC* copy number distributions of xenografts increased significantly, with a marked increase following EP (**Fig. 6D-E**). This demonstrated a selective advantage for cells with high ec*MYC* in the presence of these drugs. Together with the low number of γH2AX foci in MYC^high^ cells during treatment, these results suggest that ec*MYC* confers dose-dependent resistance to DNA-damaging agents.

## DISCUSSION

For most solid tumors, including NSCLC, chemotherapy is contraindicated if cancer progression has incapacitated a patient to the point of hospitalization. Chemotherapy acts too slowly to reverse advanced disease before the side effects exact a toll that the patient cannot endure. This rule is broken for SCLC because first-line chemotherapy works so reliably and quickly that the benefit outweighs and outpaces the side effects (49). Critically ill patients can recover to leave the hospital after one cycle of treatment. Equally reliable is that this response is transient, and after relapse, the benefit of additional lines of therapy diminishes rapidly.

One of the enduring mysteries in SCLC is that when it acquires resistance to one DNA-damaging regimen, it usually acquires cross-resistance to others. This aspect of the disease has proven stubbornly difficult to recapitulate in the laboratory, and the molecular alterations underlying this transformation remain unknown. Here we reported 51 SCLC PDX models with comprehensive genomic, transcriptomic, functional, and clinical annotation. For individual patients, serial PDXs derived before therapy and after acquired resistance reproduced this transition. For population-level characteristics of acquired resistance, the full panel of PDX models captured the distinction between untreated and relapsed SCLC, and among relapsed cases, the distinctions between platinum-sensitive and platinum-resistant cancers (CTFI < 90 days), and between SCLC that continues to respond to therapy and SCLC that does not respond. These models constitute an experimental system that captures clinical cross-resistance in SCLC with high fidelity.

We used this system to test recurrence of candidate drivers of cross-resistance. To discover these candidates, we performed an integrated genomic comparison of serial models derived from patient MGH1518. Following second-line OT, this patient’s SCLC acquired cross-resistance, and during OT, it acquired an extrachromosomal DNA amplification of *MYC* (ec*MYC*) that radically altered the transcriptome. We exploited the intrinsic random segregation of ecDNAs to demonstrate that ec*MYC* was a dose-dependent driver of cross-resistance. Of note, this approach can be applied broadly to assess the impact of ec*MYC* on cancer cell fitness under a variety of experimental conditions. Across the PDX panel, we found an additional cross-resistant PDX with ec*MYC* (MGH1564-1A) and a second serial pair in which ec*MYCN* was acquired after therapy (MGH1578). Overall, ecDNA amplifications of *MYC* paralogs were enriched in cross-resistant models from relapsed patients.

The current consensus on the molecular evolution of SCLC during disease progression is that genetic alterations drive initial tumorigenesis and possibly metastasis, but thereafter epigenetic mechanisms drive resistance to therapy (8). Promising candidates include upregulation of multi-drug-resistance efflux pumps (50), soluble guanylate cyclase (sGC) (51), the Wnt signaling pathway (52), and the epithelial-to-mesenchymal transition (53,54), as well as downregulation of the putative RNA/DNA helicase SLFN11 (54-57) and neuroendocrine differentiation factors (58). However, discovering the initiating events that induce these gene expression changes has proven challenging.

To the best of our knowledge, ec*MYC* is the second genomic driver of resistance to be discovered in an SCLC patient for any therapy, and the first acquired driver of cross-resistance. The previous driver of resistance was discovered in 1983, also on an ecDNA (59). An SCLC patient was treated for 12 months with high-dose methotrexate (MTX), and at progression, the cell line NCI-H249P was derived in the presence of MTX. This cell line harbored an ecDNA amplification of dihydrofolate reductase (ec*DHFR*), the target of MTX. Unlike ec*MYC*, ec*DHFR* is specific for MTX resistance, and retention in NCI-H249P required continuous MTX treatment. MTX has since fallen out of the SCLC armamentarium, but the ec*DHFR* example raises the possibility that there may be other ecDNAs with drug-specific resistance genes that lack a drug-independent selective advantage, and these may vanish during model derivation unless drug treatment is maintained. Both ecDNAs and *MYC* paralog amplifications have a 40-year history of research in SCLC. Amplification of *MYC* was the first genomic alteration to be identified in SCLC (also in 1983 (60)), followed by *MYCL* (61) and *MYCN* (62), preceding the discoveries of biallelic loss of *RB1* and *TP53* that became the genomic hallmarks of this disease (63,64). Cytogenetic analyses revealed frequent *MYC* paralog amplifications on ecDNAs (double-minute chromosomes) (60,62), and these were corroborated by genome sequencing of an SCLC cell line as one of the first examples of chromothripsis (65). The pathologic functions of these genes have been continuously investigated in SCLC, with roles described in tumorigenesis, metastasis, intratumoral heterogeneity, and lineage plasticity (66-71). However, the ec*MYC* amplification in MGH1518-3A played no role in initial tumorigenesis or metastasis. It was acquired during therapy and conferred drug resistance, and the roles of *MYC* paralogs in drug resistance are less clear.

The largest WGS profile of human SCLC tumors revealed recurrent amplifications of *MYC, MYCN*, and *MYCL* (6%, 4% and 9%, respectively), and a recent AmpliconArchitect analysis of this data revealed that most were on ecDNAs (44,72). However, these were primarily surgical resection specimens, which strongly tilted the analysis towards early-stage patients (80% of samples) who had not received chemotherapy (95% of samples) (72). The poor prognosis observed among ecDNA+ patients may signify the likelihood of relapse after surgery, but functional testing of these fixed chemo-naïve specimens for cross-resistance was not possible (44). The cell lines established at the NCI-Navy Medical Oncology Branch from 1977-1992 may be more relevant for acquired cross-resistance, as most were from patients with advanced SCLC who received chemotherapy, and approximately half were established post-relapse (73). Amplifications of *MYC* paralogs were significantly more frequent in cell lines derived after relapse, suggesting a role in acquired resistance (74-76). More recently, ectopic expression experiments in a chemo-sensitive PDX derived from an untreated SCLC patient demonstrated that both MYCN and MYCL can confer chemo-resistance (77). The main advance in this work is to develop models that recapitulate clinical cross-resistance, among them isogenic serial models that directly demonstrate the resistance conferred by acquired ec*MYC* amplification.

Our findings suggest that, for many patients, ec*MYC* amplifications transform SCLC into an untreatable disease. This discovery opens major epidemiologic, mechanistic, and therapeutic questions. The fraction of refractory SCLC cases that can be ascribed to ec*MYC/N/L*, and whether each paralog contributes equally, remains to be determined. Non-invasive assays such as FISH on interphase CTCs may enable prospective monitoring of ecDNAs to answer these questions. The mechanism by which ec*MYC* causes resistance is largely unexplored, including whether the resistance functions are distinct from those that drive tumorigenesis, and whether complete elimination of ec*MYC* is necessary to re-sensitize. Pharmacologic targeting of MYC is one of the enduring challenges in cancer research, but strategies to disrupt ecDNA maintenance mechanisms could also be effective. Finally, the discovery of one category of cross-resistance may help to distinguish others and to determine whether they are overlapping or mutually exclusive. A landscape of cross-resistance drivers would be a major step towards personalized therapy for relapsed SCLC.

## METHODS

### PDX model generation

All mouse studies were conducted through Institutional Animal Care and Use Committee–approved animal protocols in accordance with Massachusetts General Hospital and University of Texas Southwestern Medical Center institutional guidelines. PDX models were generated from CTCs, biopsy/resection specimens, or malignant effusions as described previously (35). The following 9 PDX models are first reported in this study: MGH1505-1A, MGH1508-1A, MGH1517-1A, MGH1567-1A, MGH1572-4A, MGH1578-1A, MGH1709-2A, MGH1717-1A, MGH1719-1A. All other models have been reported previously (35,36,78). Palpable tumors were measured with electronic calipers until tumor volume exceeded 1500 mm^3^. Measurement frequency was weekly unless drug treatment was initiated. Tumor volume was estimated by using the spheroid formula: tumor volume = [(tumor length) × (tumor width^2^)] × 0.52. Upon mouse euthanasia and xenograft resection, scalpel-dissected fragments were either implanted immediately into the right flank of an NSG mouse (NOD.Cg-Prkdc^scid^ Il2rg^tm1Wjl^/SzJ) for passaging, processed for downstream molecular analysis or short-term culture growth, or preserved in cryopreservation media for future use. For downstream molecular analyses xenograft tissue samples were either fixed or flash-frozen within 5 minutes of mouse euthanasia. Tissue samples were fixed overnight in 10% neutral buffered formalin for histology and immunohistochemistry studies, or fresh-frozen in liquid nitrogen for western blot and nucleic acid assays.

### PDX model drug testing

Drugging studies were initiated at xenograft volumes = 300-600 mm^3^ and tumors were measured twice weekly at least three days apart. Drugs were used as follows (unless otherwise stated): EP treatment cycle – cisplatin (7 mg/kg, intraperitoneal) once on day 1, and 8, and etoposide (10 mg/kg, intraperitoneal) once on day 1-3 and 8-10. OT treatment cycle - Olaparib (50 mg/kg, oral gavage) twice daily for 5 days, TMZ (25 mg/kg, oral gavage) for 5 days once daily. Drugging with TMZ was performed 4 hours after Olaparib treatment, and a gap of 8-9 hours was maintained between two Olaparib administrations. Topotecan (1 mg/kg, intraperitoneal) was administered once daily on day 1-5 and 8-12. For analysis, tumor measurement was included for tumor volume > 2× ITV or 50 days after start of treatment, whichever was attained earlier.

### PDX model drug response analysis

PDX drug responses were quantified as a function of change in tumor volume with or without treatment over time, as depicted for MGH1518-1BX treated with EP in Supplementary Figure S1A-C. Tumor volume was expressed relative to initial tumor volume (ITV) on first day of treatment, within endpoints of 50 days or 200% ITV. For treated xenografts, area under the tumor volume curve (AUC) was estimated as the sum of right trapezoids between volume measurements, bounded by endpoints. The AUC for a model∼regimen combination (AUC^drug^) was calculated as the mean AUCs of all xenografts treated with that regimen. In the absence of treatment, some PDX models grew too rapidly for precise measurement of tumor doubling (200% ITV) using twice-weekly caliper measurements. Therefore, the exponential growth rates of untreated xenografts were calculated by linear regression of log-transformed volume measurements during growth from a starting range of 100-300 mm^3^ to a final range of 1200-1500 mm^3^. The mean growth constant for each model was calculated from 6-94 replicate xenografts (median 16 per model) and used to estimate the AUC without treatment to 200% ITV over 50 days (AUC^no.drug^). The effect of a drug regimen on model tumor volume was calculated as the difference between AUC^no.drug^ and AUC^drug^, ΔAUC. All volume calculations, tumor volume curves and derived metrics were generated in R from compiled spreadsheets containing xenograft identities and bi-weekly electronic caliper measurements.

### PDX short-term cultures (STC)

Xenografts were resected and collected in cold PBS and the cells were manually dissociated using a scalpel. Live cell clusters were enriched by repeated passes of gravity sedimentation in 15 ml conical tubes, with interval assessment of supernatant to identify fractions with live cell clusters and minimal debris. ACK lysis buffer (Gibco, Cat # A1049201) was used to remove red blood cells. Cells clusters were plated and grown in HITES media with 5% FBS until use.

### Whole genome sequencing

Genomic DNA was extracted from flash-frozen PDX tumor tissue and germline controls using the Qiagen DNeasy Blood & Tissue Kit (Cat# 69506). Whole-genome sequencing of the 51 PDX models (41 with matched germline control) was performed by Novogene (www.novogene.com; NovaSeq 6000 PE150 with 60X coverage). Sequencing reads were aligned to the human reference genome GRCh38.v21 or the mouse reference genome GRCm38.p3 using bwa-0.7.12 (github.com/lh3/bwa). Mouse reads were filtered when the mouse alignment score (AS flag in bam files) was greater than the human alignment score. Duplicated reads were then marked using Picard MarkDuplicates (v2.27.5; https://broadinstitute.github.io/picard/).

### Somatic variants

Somatically acquired variants in the 41 germline-matched samples were determined with Mutect2 and FilterMutectCalls from GATK (gatk-4.1.2; github.com/broadinstitute/gatk). For the 10 non-matched samples, we used HaplotypeCaller, SelectVariants, and VariantFiltration from GATK. Gene annotation was then done for both matched and unmatched samples with SnpEff (pcingola.github.io/SnpEff/). Potential germline variants in unmatched samples were removed using the NCBI dbSNP database of common germline variants: (ftp.ncbi.nih.gov/snp/organisms/human_9606_b151_GRCh38p7/VCF/common_all_20180418.vcf.gz, minus variants from COSMIC [cancer.sanger.ac.uk/cosmic]) as well as our own set of 187 germline controls. Further variant filtering was done by removing variants with allele frequency < 0.01 and allele depth < 2, by removing variants with LOW impact (in SnpEff output).

### Copy number variants

Copy number variation (CNV) was estimated from the distribution of exon read depth in whole-genome sequencing using circular binary segmentation (implemented in the R package ‘DNAcopy’; Olshen et al, Biostatistics (2004), 5, 4, pp. 557–572) with the following adjustments: *Choice of diploid controls*: Because of possible batch effects, diploid controls were generally selected from the same sequencing run as tumor samples (but not necessarily from the same patient DNA). *Recalibration*: Because tumor samples are generally not diploid overall, copy numbers were recalibrated by visual inspection of each DNAcopy-generated plot so that areas of the genome with 2 copies should have log2(tumor/control) = 0. Two CNV outputs were obtained, one with “raw” exon copy numbers that are highly variable and one with “segmented” copy numbers that contains large areas of constant copy number. Since the latter tends to obscure small focal amplifications, these were put back by searching the raw CNV data for areas of at least 3 highly amplified consecutive exons.

### Amplicon structural analysis

For amplicon analysis, aligned WGS bam files were processed by AmpliconSuite-pipeline (v0.1477.1, previously named as PrepareAA) to generate seeds for AmpliconArchitect (v1.3.r3), which returned the architectures of amplicons. Then, outputs of AmpliconArchitect were interpreted by AmpliconClassifier (v0.4.16) to annotate the types of amplicons (ecDNA, breakage-fusion-bridge, complex non-cyclic, and linear amplification). Finally, circular plots of ecDNAs were generated by CycleViz (v0.1.3).

### Mutation signature analysis

Mutations called with Mutect2 were filtered by a variant allele frequency of greater than or equal to 0.01 and those with a VAD greater than or equal to 2. To deconvolve mutation signatures, single-nucleotide substitutions from 39 SCLC whole genome sequences with matched normal were tallied by their 3’ and 5’ reference base resulting in a 96-category base substitution matrix. Using nonnegative matrix factorization (k=4), mutational signatures were calculated in MATLAB (v2017b). Mutational signatures are displayed based on the trinucleotide context and substitution type, with the frequency of mutations represented by the bar height (“Lego” plot). The four major deconvolved mutational signatures were those associated with smoking, APOBEC, aging, and TMZ. MGH1518-1BX showed a mutational pattern associated with smoking, SBS4, dominated by C>A mutations regardless of 3’ or 5’ context. MGH1518-3A showed a striking pattern of numerous C:G > T:A mutations, SBS11, associated with temozolomide methylation of guanines. To determine the variant allele frequency distribution across the identified *MYC*-containing ecDNA, mutations were filtered to single-nucleotide variants representing the associated TMZ substitutions C:G > T:A and restricted to those within the regions identified by AmpliconArchitect as part of the complex ecDNA.

### RNA sequencing alignment, mutation calling and transcript abundance measurement

RNA was extracted from flash frozen PDX tumor tissue using the Qiagen RNeasy Mini Kit (Cat# 74104). RNA quality assessment (1% agarose gel electrophoresis, Nano-drop for RNA amount and purity, Agilent2100 for RNA integrity number), library preparation (Poly-A enrichment: 250-300 bp insert cDNA library), and paired-end RNA sequencing (NovaSeq PE150 strategy) were performed by Novogene or as previously described (35,36). Samples with an RNA integrity number (RIN) of > 8 were selected for library construction. Sequencing reads were aligned to the human reference genome GRCh38.v21 or the mouse reference genome GRCm38.p3 using STAR-2.7.1a (github.com/alexdobin/STAR). Mouse reads were filtered when the mouse alignment score (AS flag in bam files) was greater than the human alignment score. Duplicated reads were then removed using Picard MarkDuplicates (v2.27.5; broadinstitute.github.io/picard/). FPKM values were generated with cufflinks-2.2.1 (cole-trapnell-lab.github.io/cufflinks/) and then normalized to transcripts per million (TPM). Alternatively, mouse-filtered bam files were reverted to fastq format with samtools fastq (samtools-1.16; www.htslib.org) then used to generate TPMs with salmon-1.9.0 (github.com/COMBINE-lab/salmon) and R package tximport (bioconductor.org/packages/release/bioc/html/tximport.html).

### Chromatin immunoprecipitation sequencing (ChIP-Seq)

Nuclei and cross-linked chromatin were prepared from flash-frozen xenograft tissue as previously described (79). Briefly, fragments were cut by scalpel, resuspended in buffer containing 0.1% NP40, 0.1% Tween-20 and 0.01% Digitonin, homogenized using a TissueLyser II (Qiagen) set at 30 Hz for 1 minutes and incubated on ice for 10 minutes. Lysates were filtered through a 40 μm strainer. Nuclei collected after cold centrifugation were cross-linked with 1.5% formaldehyde for 10 minutes and quenched with 0.125 M glycine. Sonication was performed on a Bioruptor (Diagenode) on high power, 30 sec on/off (50/50) for 25 minutes at 4°C, and samples were centrifuged 20,000 g x 10 min at 4°C. An aliquot of supernatant was frozen for chromatin input control. Chromatin was immunoprecipitated overnight with H3K27ac antibody (Abcam cat# ab4729) conjugated to Dynabeads (LifeTech). Samples were eluted at 65°C in 50 mM Tris, 10 mM EDTA, 1% SDS and cross-links were reversed overnight. DNA was extracted using MaXtract High Density tubes (Qiagen) and ChIP-seq libraries were prepared using the ThruPLEX DNA-Seq Kit (Takara). 75 bp single-end reads were sequenced on a Nextseq instrument (Illumina). Reads were aligned by bwa-mem2 (v2.2.1) to hg38 reference genome. Reads were then sorted, and duplicates were removed, by sambamba (v1.0.0).

### Western blotting

Whole cell protein was extracted from flash frozen PDX tumor tissue. Tissue samples were lysed in RIPA buffer supplemented with protease and phosphatase inhibitors (Roche, Cat# 04693159001, 04906837001) using TissueLyser II (Qiagen) homogenizer at 4°C following the manufacturer’s instructions. Samples were then centrifuged for 10 min at 15000 g and supernatant was collected for the assay. Protein concentration was measured by the Pierce BCA protein assay (ThermoFisher, Cat #23225). Samples were separated on 4-20% Mini-PROTEAN® TGX™ Precast Protein Gels (Bio-Rad) after heat denaturation in Laemmli sample buffer, and proteins were transferred to nitrocellulose membranes. Membranes were blocked with EveryBlot Blocking Buffer (Bio-Rad, Cat# 12010020) for 10 min at room temperature. The membrane was probed overnight at 4°C with the following antibodies: Myc (1:1000; Cell Signaling, 5605S), Cyclophillin A (1:1000, Cell Signaling, 2175s) followed by incubation with secondary antibody (Li-Cor, IRDye 800CW Goat anti-Rabbit IgG (1:15000) and IRDye® 680RD Goat anti-Mouse IgG (1:15000)) in EveryBlot Blocking Buffer for 1 hr at room temperature. Following washes in TBST, membranes were imaged using Rio-Rad ChemiDoc MP imaging system.

### Tissue fixation and sectioning

Freshly dissected xenograft samples were fixed overnight at 4°C in 10% neutral buffered formalin, rinsed in PBS 3 × 5 min on ice and stored in 70% ethanol for up to 1 week. Dehydration and paraffin embedding were performed by standard method with an automated machine by the UT Southwestern Tissue management and Pathology Core. Formalin-fixed paraffin-embedded (FFPE) tissue blocks were incubated in room-temperature water for 2 min followed by 2 min in ice cold water before sectioning. Tissue sections were cut at 4 μm in thickness with a microtome. Sections were transferred to positively charged slides (Fischer Scientific, Cat# 1255015) and allowed to air-dry overnight before staining.

### Hematoxylin & eosin (H&E) staining, immunohistochemistry (IHC), and immunofluorescence (IF)

Slides with FFPE sections were warmed for 20-30 min in a 60°C oven before deparaffinization. The sections were then deparaffinized with xylene 3 × 5 min, rehydrated by incubation in 100% ethanol 2 × 5 min, 90 % ethanol 1 × 3 min, 70 % ethanol 2 × 3 min, 50% ethanol 1x 3 min, 30 % ethanol 1 × 3 min, and 2 × 10 min in deionized water and stained with hematoxylin and eosin (H&E). For IHC and IF, post rehydration, antigen retrieval was performed in SignalStain® Citrate Unmasking Solution (Cell Signaling, Cat# 14746S) at 98°C for 10 - 20 min. For IHC, following antigen retrieval and inhibition of endogenous peroxidase activity with 3% H2O2 for 15 min, the slides were incubated with 10% normal goat serum for 1 hr at room temperature. Tissue sections were incubated with primary antibody: Myc (1:100; Cell Signaling, 5605S) diluted in SignalStain® Antibody Diluent (Cell Signaling, Cat# 8112) for overnight at 4°C in a humid chamber. Following 3 × 5 min washes in TBST, tissue sections were incubated with HRP-conjugated secondary antibody (Vector Laboratories) for 1 hr at room temperature followed by development using chromogenic substrate DAB (SignalStain® DAB Substrate Kit, Cell Signaling, Cat# 8059s) for 10 min at room temperature. Slides were scanned and images were captured using a NanoZoomer (Hamamatsu). For IF, following antigen retrieval, and sections were permeabilized for 45 min in 0.2% Triton X in PBS. Sections were blocked in 10% goat serum, 0.05% Triton X, and 0.05% Tween 20 in PBS for 1 hr followed by incubation in primary antibodies ((MYC 1:100, Abcam Cat# 32072, γH2AX 1:100 Abcam Cat# 303656) for overnight at 4°C in a humid chamber. Following 3 × 10 min washes in PBST sections were then incubated with secondary antibodies for 60 min at room temperature. After 3 × 10 min washes in PBST, sections were stained with 4′,6-diamidino-2-phenylindole (DAPI) for 10 min and mounted using SlowFade Diamond Antifade Mountant with DAPI (Invitrogen, Cat# S36964). Slides were imaged using a Nikon Y-FL microscope attached to a Nikon DS-Qi2 camera and images were captured using NIS elements AR software. MYC intensity and the number of γH2AX foci within each nucleus were analyzed by CellProfiler (v4.2.4).

### Fluorescence in-situ hybridization (FISH)

For metaphase FISH, cultured cells were treated with 30 ng/ml KaryoMAX Colcemid overnight and washed once by PBS. Cells were then resuspended in 75 mM KCl and swollen at 37°C for 30 min. Next, cell samples were fixed by 3:1 methonal:glacial acetic acid (v/v) for 4 times and dropped onto microscopic slides. Prior to FISH probe (purchased from Empire Genomics) hybridization, slides were dried overnight, equilibrated by 2× SSC buffer, and serially dehydrated by 70%, 85%, and 100% ethanol. Slides were then denatured at 75°C for 2 min and hybridized at 37°C overnight. Finally, slides were washed twice with 0.4× SSC with 0.3% IGEPAL CA-630 and once with 2× SSC with 0.1% IGEPAL CA-630, stained with DAPI, and mounted with anti-fade media. For FISH on FFPE samples, slides were aged at 75°C for 20 min and deparaffinized by xylene. After being washed with 100% and 70% ethanol, slides were treated with 0.2N HCl at room temperature for 20 min, 10 mM citric acid at 90°C for 20 min, rinsed with 2× SSC, and digested briefly (< 1 min) by 1% Proteinase K (NEB) in TE buffer at room temperature. Next, slides were serially dehydrated with ethanol and hybridized as above. To quench autofluorescence, slides were treated with Vector TrueVIEW (Vector Laboratories) following the instructions from the manufacturer. FISH images were taken by ZEISS Apotome 3 wide-field microscope with a 63× oil lens. The total area of FISH signal within each nucleus was analyzed by CellProfiler (v4.2.4).

### Statistics

Statistical methods are indicated in the corresponding figure legends. All statistical tests are two-sided. Unless otherwise indicated in figure legends or methods, all statistical tests were performed in GraphPad Prism 9. Differential gene expression analysis was performed with R (v4.2.3) using package DESeq2. Gene set enrichment analysis was performed using the GSEA function in the clusterProfiler R package. Regression analyses were performed with R using the linear model fitting function lm().

## Acknowledgments and Funding

We are grateful to the patients and families who participated in these research studies. We thank the members of the MGH and UT Southwestern thoracic oncology groups and staff for assistance with this research. We are grateful to John D. Minna for his ongoing scientific guidance and critical reading of this manuscript. This work was supported by NCI grant U01CA220323 (N.J.D.), the Rullo Family Foundation (M.S.L.), the Cancer Prevention and Research Institute of Texas (CPRIT) grant RR210034 (S.W.), the University of Texas SPORE in Lung Cancer award number P50CA070907 (L.G.; J. Minna P.I.), CPRIT grant RR20007 (B.J.D.), NIH grant 1K08CA237832 (B.J.D.) and a Disease-Oriented Scholar Award from UT Southwestern Medical Center (B.J.D.). This work was delivered as part of the eDyNAmiC team supported by the Cancer Grand Challenges partnership funded by Cancer Research UK (P.S.M CGCATF-2021/100012, S.W. CGCATF-2021/100023) and the National Cancer Institute (P.S.M. OT2CA278688, S.W. OT2CA278683). We thank the UT Southwestern Whole Brain Microscopy Facility (RRID:SCR_017949) for assistance with whole slide scanning. Research reported in this publication was supported by the National Cancer Institute of the National Institutes of Health under award number P30CA142543.

## TABLE LEGENDS

**Table 1. Differential Gene Expression for MGH1518 Serial Models.** Gene identifiers include Ensembl gene ID, gene symbol and gene type. DeSeq2 metrics presented include baseMean, log2 Fold Change, lfcSE, stat, p.value, and p.adj. PDX transcript abundance in transcripts per million (TPM), with passage number of included xenografts indicated (psg).

**Table 2. Mutations in the ec*MYC* region.** Xenograft = PDX model and passage number on which WGS was performed. Position = chromosome 8 position of mutation (hg38). Reference = reference allele. Alternate = alternate allele. MAF = mutant (alternate) allele frequency.

**Table 3. Focused PDX model clinical, molecular, and functional profiles.** Clinical history of patient treatment with chemotherapy. Tumor source type for PDX generation. Average model AUC to 200% ITV over 50 days for untreated xenografts or xenografts treated with EP, OT or topotecan, as well as average ΔAUC of *in vivo* regimens. Presence of ecDNA in model and presence of *MYC* family member on ecDNA, and expression levels (TPM) and copy numbers of *MYC* family members.

