## Supplementary Figures for "Acquired Cross-resistance in Small Cell Lung Cancer due to Extrachromosomal DNA Amplification of *MYC* paralogs"

### Supplementary Figure S1

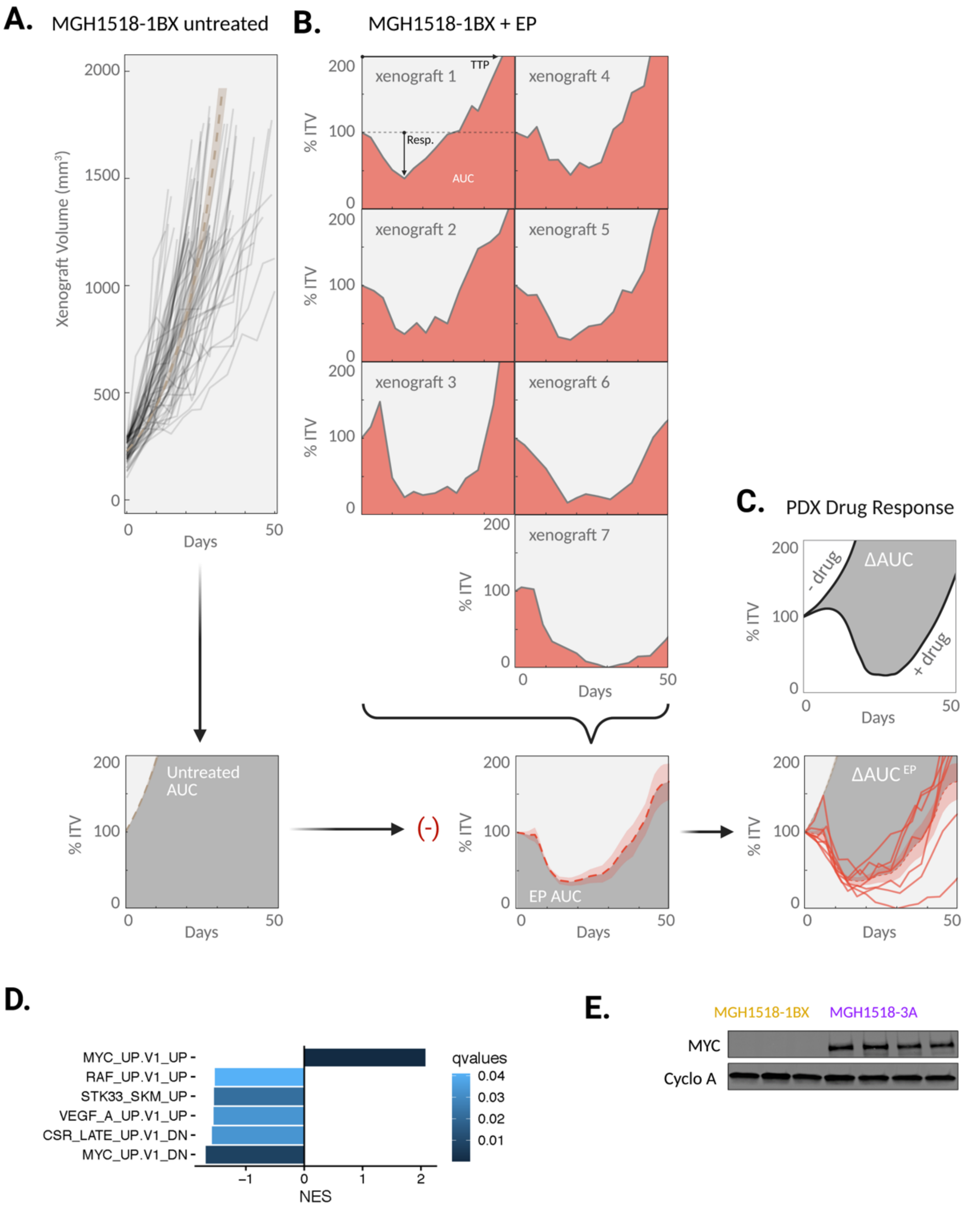


**Supplementary Figure S1.** (**A-C**) Calculation of ΔAUC for MGH1518-1BX treated with EP. (**A**) (Top) Untreated tumor-volume (TV) curves for 94 replicate xenografts of MGH1518-1BX from starting volume of 100-300 $mm^3$ to final volume of 1200-1500 $mm^3$. For each xenograft, growth coefficient was derived from linear regression of log-transformed tumor volume measurements. Tan dashed line + shading: exponential growth model of MGH1518-1BX from 94 xenografts with mean initial tumor volume (ITV) of 227.3 $mm^3$ and mean growth coefficient of 0.0659 +/- 0.0029 (SEM). (Bottom) Exponential growth model of MGH1518-1BX to volume endpoint 200% ITV over 50 days. Gray shade = area under the untreated TV curve (AUC). (**B**) (Top) TV curves for 7 replicate xenografts of MGH1518-1BX treated with EP starting at ITV = 300-600 $mm^3$, with measurements every 3-4 day until days until endpoints of 200% ITV or 50 days. Red shade = area under the TV curve (AUC) estimated from sum of right trapezoids between TV measurements. Arrows indicate metrics of maximum tumor regression (best response, “Resp.”) or time to volume endpoint (time to progression, “TTP”) for the first xenograft. (Bottom) Mean TV curve for MGH1518-1BX treated with EP +/- 95% CI. Gray shade = mean AUC for MGH1518-1BX treated with EP. (**C**). Effect of EP on MGH1518-1BX tumor volume. Gray shade = change in AUC with EP treatment ($\Delta AUC^{EP}$). (**D**) Gene set enrichment analysis of differentially expressed genes in MGH1518-3A vs. MGH1518-1BX. (**E**) MYC protein levels in replicate xenografts of MGH1518-1BX and MGH1518-3A.

### Supplementary Figure S2

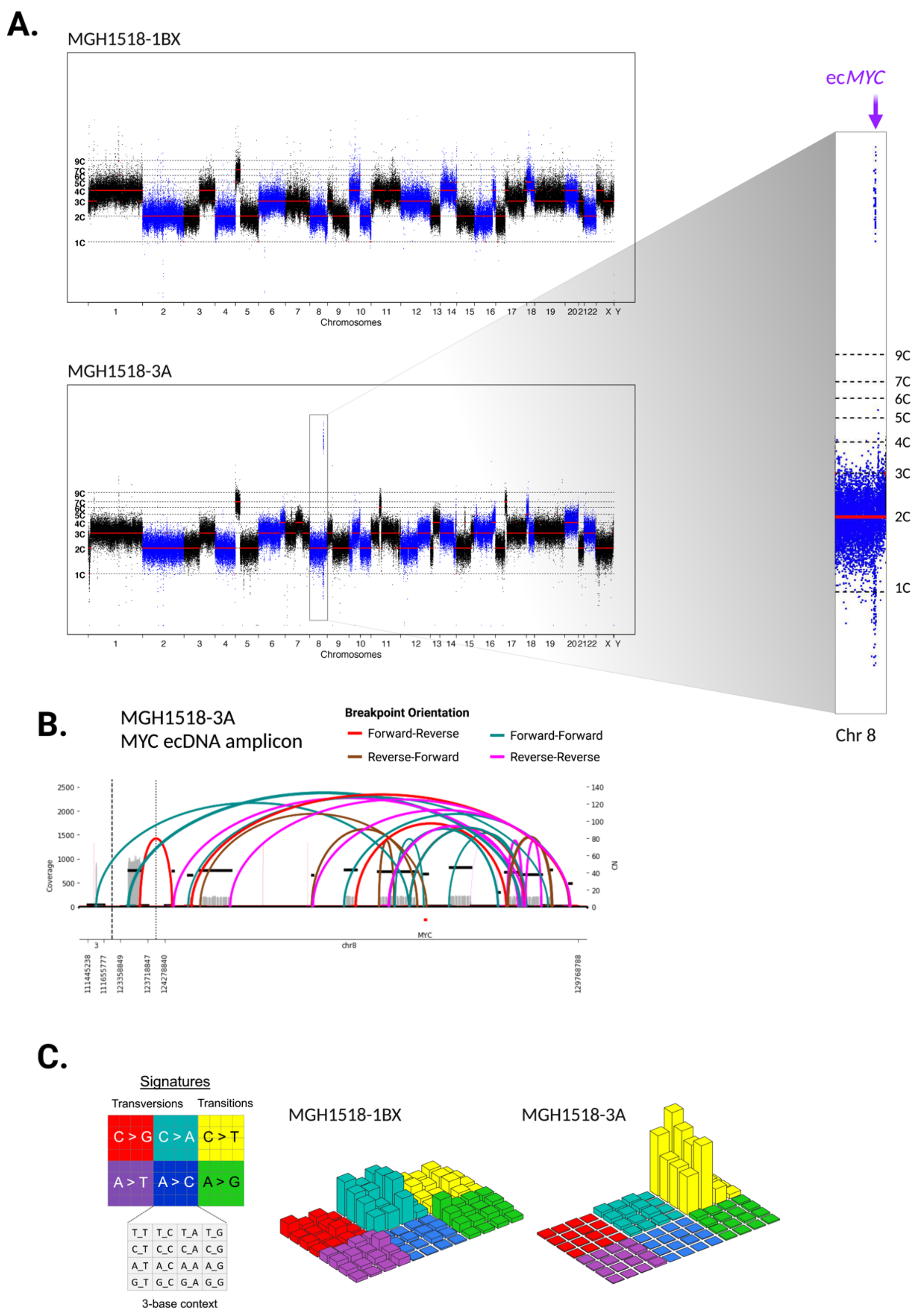


**Supplementary Figure S2.** (**A**) Copy number variation across whole genomes of MGH1518-1BX and MGH1518-3A. Inset: chromosome 8 with region of ec*MYC* amplification indicated. (**B**) AmpliconArchitect reconstruction of the rearrangements that formed ec*MYC* in MGH1518-3A. (**C**) Three-dimensional bar plots (also called “Lego plots”) representing mutational signatures in a three-base context for each model.

### Supplementary Figure S3

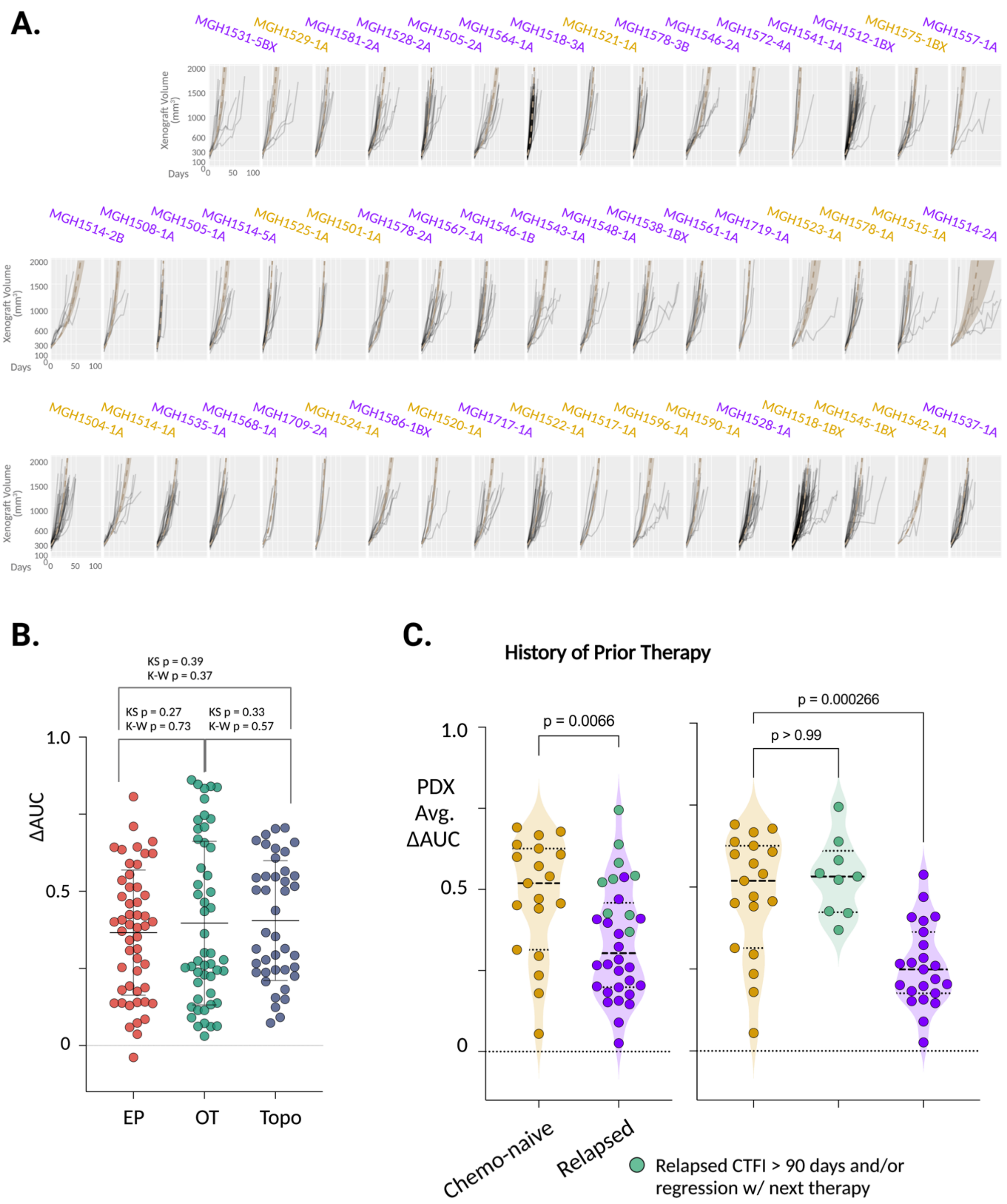


**Supplementary Figure S3.** (**A**) TV curves for untreated xenografts of each PDX model, to estimate the 50-day AUC of untreated tumor doubling, as in Supplementary Figure S1A. (**B**) Comparison of ΔAUC measurement distributions for PDX models treated with EP, OT and topotecan. KS p = two-sample Kolmogorov-Smirnov tests to determine whether paired distributions are significantly different. K-W p = Kruskal-Wallis test to determine probability that differences in the three distributions could be due to random sampling. (**C**) Comparison of $\Delta AUC^{avg}$ from PDX models derived from patients with untreated vs relapsed SCLC. Relapsed PDXs are further annotated for clinical features associated with preserved chemosensitivity (teal): derivation after first-line therapy from patients with “platinum-sensitive” SCLC (CTFI > 90 days), or derivation after ≥1 line of therapy from patients who responded to their next line of chemotherapy. (Left) Mann-Whitney test. (Right) Kruskall-Wallis test for comparison of post-relapse columns with chemo-naïve control column.

### Supplementary Figure S4

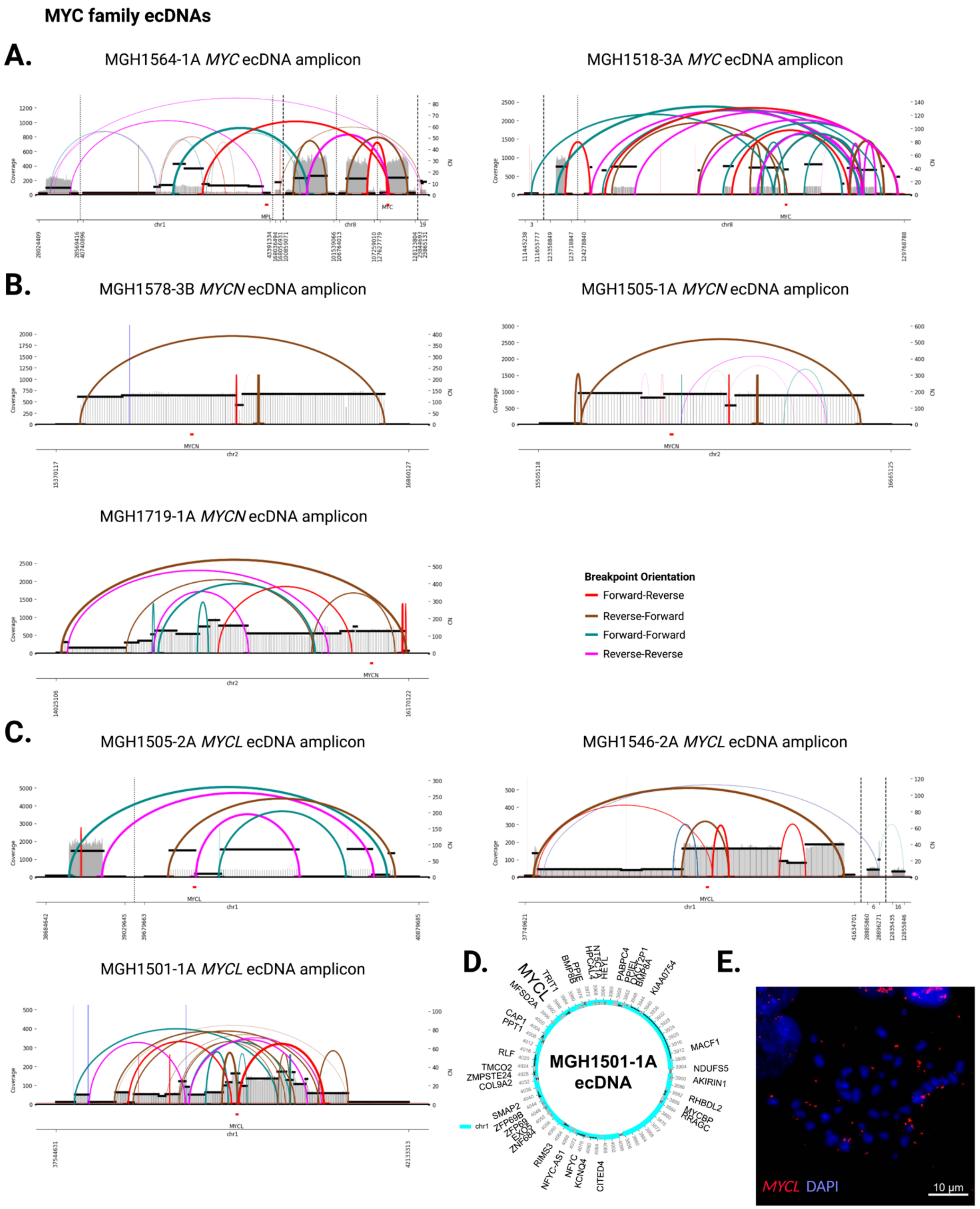


**Supplementary Figure S4.** (**A-C**) AmpliconArchitect reconstruction of the rearrangements that formed ecDNAs harboring *MYC* (**A**) or paralogs *MYCN* (**B**) or *MYCL* (**C**) in the PDX panel. (**D**) Consensus circular map of ec*MYCL* in MGH1501-1A. (**E**) FISH of probes for *MYCL* in metaphase cells from MGH1501-1A.

### Supplementary Figure S5

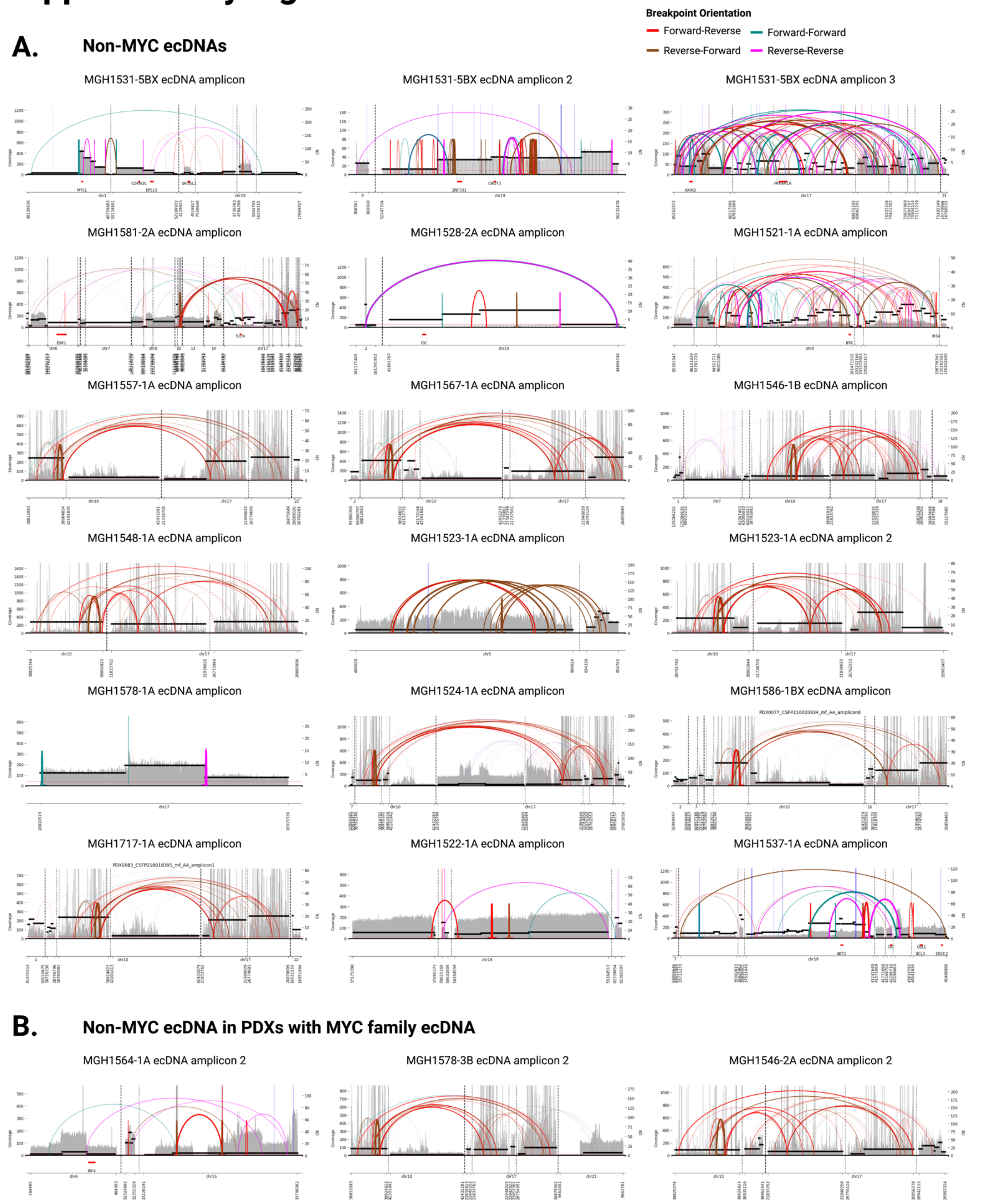


**Supplementary Figure S5.** AmpliconArchitect reconstruction of the rearrangements that formed ecDNAs that do not harbor *MYC* paralogs, either in PDXs that contain no *MYC* paralog ecDNAs (**A**) or in PDXs that also contain a *MYC* paralog ecDNA (**B**).

### Supplementary Figure S6

A.

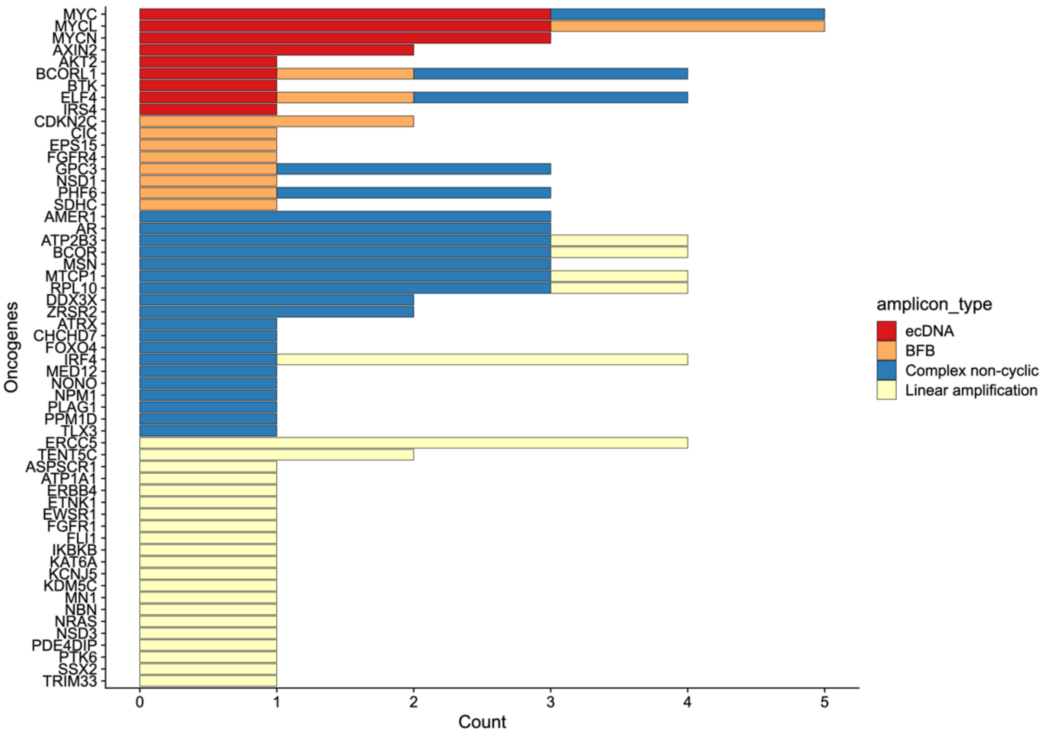


B.

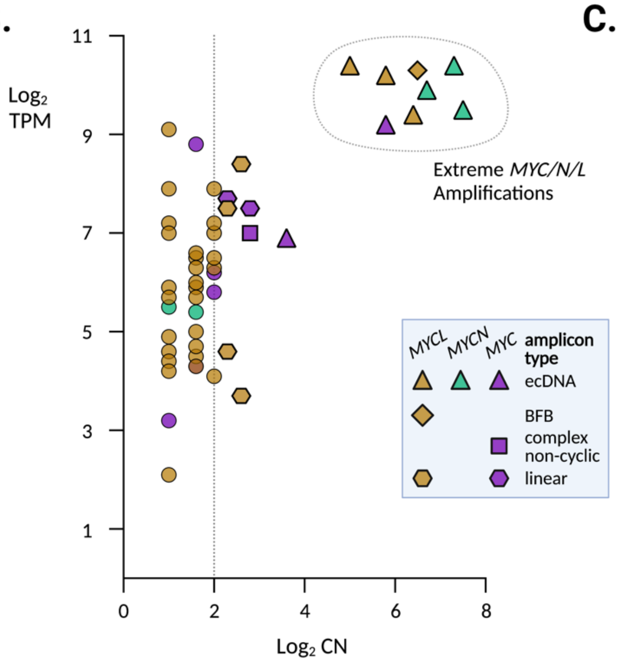


C.

| Model | Residual Standard Error | Adjusted R-squared | P-value |
|---|---|---|---|
| Delta AUC ~ MYCs DNA CN | 0.1717 | 0.1692 | 0.00910 *** |
| Delta AUC ~ MYCs mRNA | 0.1747 | 0.1340 | 0.02064 * |
| Delta AUC ~ ecMYCs | 0.1771 | 0.1098 | 0.01007 * |
| Delta AUC ~ ecDNAs | 0.1820 | 0.0601 | 0.04594 * |

| Model | Residual Standard Error | Adjusted R-squared | P-value |
|---|---|---|---|
| Delta AUC EP ~ MYCs DNA CN | 0.1888 | 0.1359 | 0.02101 * |
| Delta AUC EP ~ MYCs mRNA | 0.1888 | 0.1346 | 0.02032 * |
| Delta AUC EP ~ ecMYCs | 0.1918 | 0.1067 | 0.01106 * |
| Delta AUC EP ~ ecDNAs | 0.1960 | 0.0673 | 0.03678 * |

| Model | Residual Standard Error | Adjusted R-squared | P-value |
|---|---|---|---|
| Delta AUC OT ~ MYCs DNA CN | 0.2530 | 0.1047 | 0.04669 * |
| Delta AUC OT ~ MYCs mRNA | 0.2560 | 0.0683 | 0.10110 |
| Delta AUC OT ~ ecMYCs | 0.2562 | 0.0670 | 0.03865 * |
| Delta AUC OT ~ ecDNAs | 0.2552 | 0.0745 | 0.03092 * |

| Model | Residual Standard Error | Adjusted R-squared | P-value |
|---|---|---|---|
| Delta AUC TOPO ~ MYCs DNA CN | 0.1897 | 0.0462 | 0.19540 |
| Delta AUC TOPO ~ MYCs mRNA | 0.1877 | 0.0661 | 0.13940 |
| Delta AUC TOPO ~ ecMYCs | 0.1865 | 0.0781 | 0.04262 * |
| Delta AUC TOPO ~ ecDNAs | 0.1940 | 0.0023 | 0.30240 |

**Supplementary Figure S6.** (**A**) Oncogenes amplified in PDX models, ordered by frequency of amplification and by type of amplicon as annotated by AmpliconArchitect. (**B**) Comparison of transcript level vs. copy number for the *MYC* paralog with the highest expression level in each model. Amplicon type is annotated. Dashed vertical line at CN = 4, beyond which we classify *MYC* paralogs as amplified. A clear break is observable between models with < 15 *MYC* paralog copies and models with > 30 copies ("extreme amplifications", dashed circle). (**C**) Linear regression analysis of PDX responses to individual agent EP, OT and topotecan, as well as average drug response ($\Delta AUC^{avg}$), versus *MYC* paralog copy number, transcript level, or incorporation into an ecDNA, or presence of any ecDNA in the model.

### Supplementary Figure S7

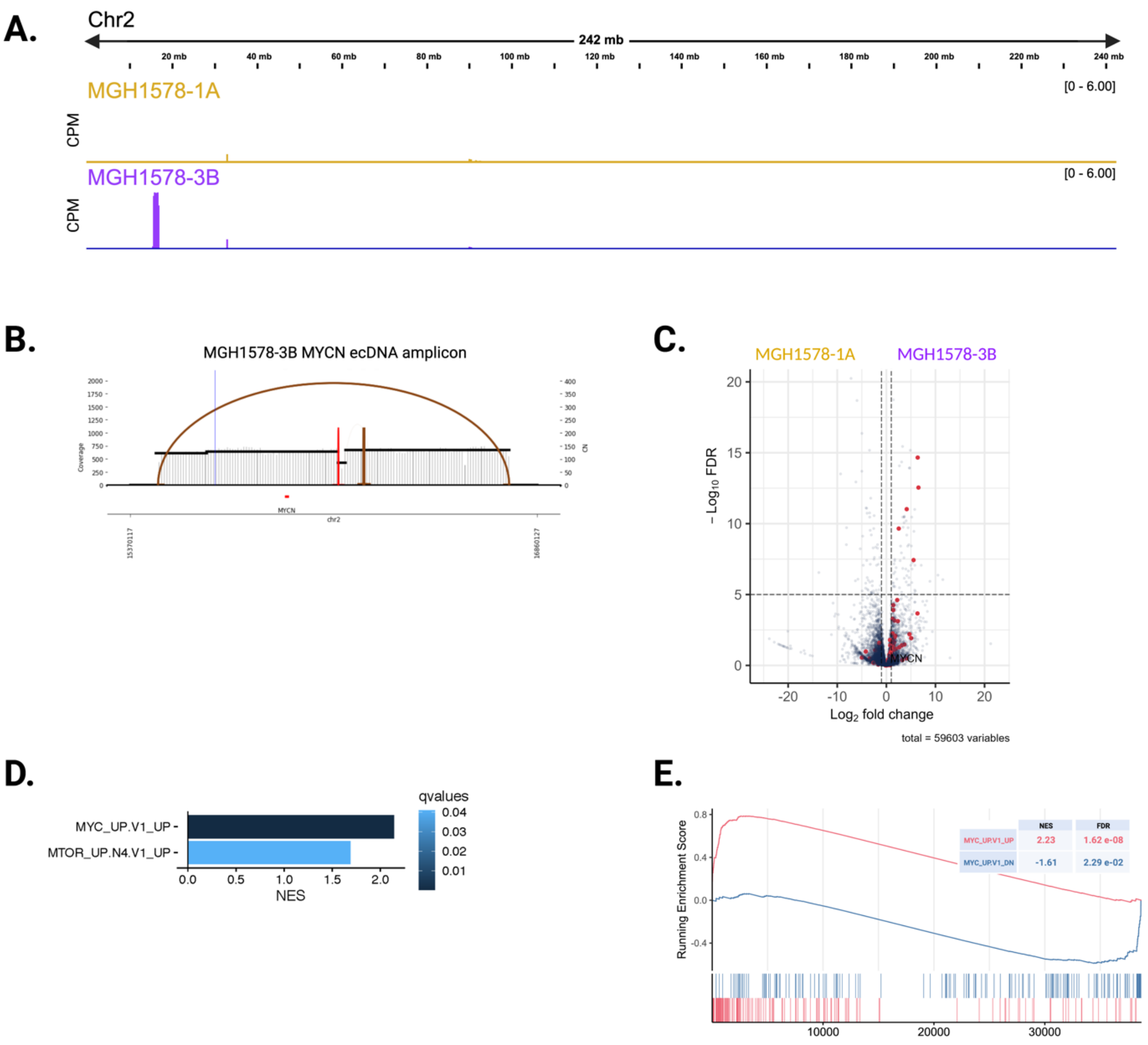


**Supplementary Figure S7.** (**A**) Focal amplification of *MYCN* on chromosome 2 in MGH1578-3B but not MGH1578-1A. Peak heights = counts per million mapped reads (CPM). (**B**) AmpliconArchitect reconstruction of rearrangements to form ec*MYCN* in MGH1578-3B. (**C**) MGH1578 PDX differential gene expression. Red = genes upregulated upon overexpression of *MYC* in primary breast epithelial cells (MYC_UP.V1_UP) as in **Figure 1E**. (**D**) Gene set enrichment analysis of differentially expressed genes in MGH1578-3B vs. MGH1578-1A. (**E**) Gene set enrichment plot in MGH1578-3B vs. MGH1578-1A for genes upregulated and downregulated with MYC overexpression, as in **Figure 1F**.
